# The killing of human gut commensal *E. coli* ED1a by tetracycline is associated with severe ribosome dysfunction

**DOI:** 10.1101/2023.07.06.546847

**Authors:** Iskander Khusainov, Natalie Romanov, Camille Goemans, Beata Turoňová, Christian E. Zimmerli, Sonja Welsch, Julian D. Langer, Athanasios Typas, Martin Beck

## Abstract

Ribosomes translate the genetic code into proteins. Recent technical advances have facilitated in situ structural analyses of ribosome functional states inside eukaryotic cells and the minimal bacterium Mycoplasma. However, such analyses of Gram-negative bacteria are lacking, despite their ribosomes being major antimicrobial drug targets. Here we compare two *E. coli* strains, a lab *E. coli* K-12 and human gut isolate *E. coli* ED1a, for which tetracycline exhibits bacteriostatic and bactericidal action, respectively. The *in situ* ribosome structures upon tetracycline treatment show a virtually identical drug binding-site in both strains, yet the distribution of ribosomal complexes clearly differs. While K-12 retains ribosomes in a translation competent state, tRNAs are lost in the vast majority of ED1a ribosomes. A differential response is also reflected in proteome-wide abundance and thermal stability assessment. Our study underlines the need to include molecular analyses and to consider gut bacteria when addressing antibiotic mode of action.

HIGHLIGHTS
• Ribosome structures of gram-negative bacteria are analyzed in situ
• Tetracyline is bactericidal to gut isolate despite identical ribosome structures
• When antibiotic is bacteriostatic, ribosomal translation competent states are retained
• When antibiotic is bactericidal, cells rapidly accumulate P-tRNAs-deficient ribosomes

**GRAPHICAL ABSTRACT**

## INTRODUCTION

Ribosomes are essential molecular machines responsible for protein synthesis by decoding genetic information from messenger RNA (mRNA) into polypeptide chains. This process requires the assistance of several translation factors and consists of four main stages: initiation, elongation, termination, and recycling. The actual synthesis of new polypeptide chains happens during repetitive rounds of elongation. During this process in bacteria, translation factor EF-Tu delivers aminoacyl-tRNA to the ribosomal A-site, ensuring its proper accommodation on the ribosome; while translation factor EF-G facilitates the translocation of the tRNAs from the A- and P-sites to P- and E-sites respectively, and promotes progression of the ribosome along the mRNA (Voorhees and Ramakrishnan, 2013). The P-site tRNA is a critical component of the translation machinery, as it carries the peptide chain and interacts with the ribosome to ensure its correct positioning for the next round of elongation. Recent advances of the in situ cryo-electron tomography (cryo-ET) expanded structural analysis of the ribosomes from in vitro reconstituted complexes with tRNAs and translation factors to visualization of their action in close to native environment and under stress in cells (Fedry et al., 2023; Gemmer et al., 2023; Hoffmann et al., 2022; Xing et al., 2023; Xue et al., 2022). Surprisingly, bacteria remain underrepresented among organisms for which translation states have been characterized at high resolution by cryo-ET and subtomogram averaging (STA). Until now, such studies have been mainly conducted on the minimal bacterium *Mycoplasma pneumonia*, and uncovered that translation inhibitors shift the distribution of ribosome states inside the cells (O’Reilly et al., 2020; Xue et al., 2022). Nonetheless, antibiotic mode-of-action, secondary targets and resistance can differ substantially across bacterial species, and sometimes strains.

Strikingly, bacteriostatic ribosome-binding antibiotics such as tetracyclines are bactericidal to some members of the gut microbiota, and may thus cause acute damage to this complex community of microorganisms (Maier et al., 2021). This means that antibiotics known to only stop bacterial growth of the few model strains tested before, also effectively kill certain gut bacteria (Maier et al., 2021). One remarkable example is the gut isolate *E. coli* ED1a (Clermont et al., 2008), which was found to be killed by tetracyclines, in particular doxycycline (Maier et al., 2021), unlike *E. coli* BW25113 (*E. coli* K-12), one of the best-characterized organisms in molecular biology (Baba et al., 2006). Although doxycycline has the same minimal inhibitory concentration (MIC) of 4 µg/ml for both strains (Maier et al., 2021), they apparently respond to the treatment differently.

It remains unclear why a gut commensal *E. coli* strain would show such a distinct reaction to a drug compared to the lab strain, especially when treated with well-described and established antibiotics such as tetracylines. The primary target of tetracycline (hereafter referred to as TET) is the A-tRNA binding site of the small subunit of the bacterial ribosome (Brodersen et al., 2000), and its bacteriostatic action is mainly attributed to its reversible binding. Structures of the 70S ribosome in complex with TET solved for gram-negative bacteria *T. thermophilus* and *E. coli* SQ171 (Cocozaki et al., 2016; Jenner et al., 2013), together with their structure alignment to a vacant ribosome of gram-positive *Staphylococcus aureus* (Khusainov et al., 2016), suggest that the TET binding mode is overall conserved.

Despite the known primary ribosomal binding site of TET, it is still unknown to which extent TET impacts the translation apparatus inside cells. Specifically, it is unclear if TET shifts translation states, whether ribosomes remain as intact 70S particles, and/or if a pool of ribosomes is potentially protected in a hibernation-like manner (Prossliner et al., 2018; Yoshida and Wada, 2014). Understanding the ribosome dynamics would thus allow further characterization of TET’s mode of action in situ.

Here, we apply a number of orthogonal methods, including thermal stability quantitative proteomics, *in situ* cryo-electron tomography (cryo-ET), subtomogram averaging (STA), and single particle cryo-electron microscopy (**Figure 1**) to investigate how the translation system copes with exposure to TET in the lab strain *E. coli* K-12 in comparison to the gut isolate *E. coli* ED1a.

**Figure 1.**
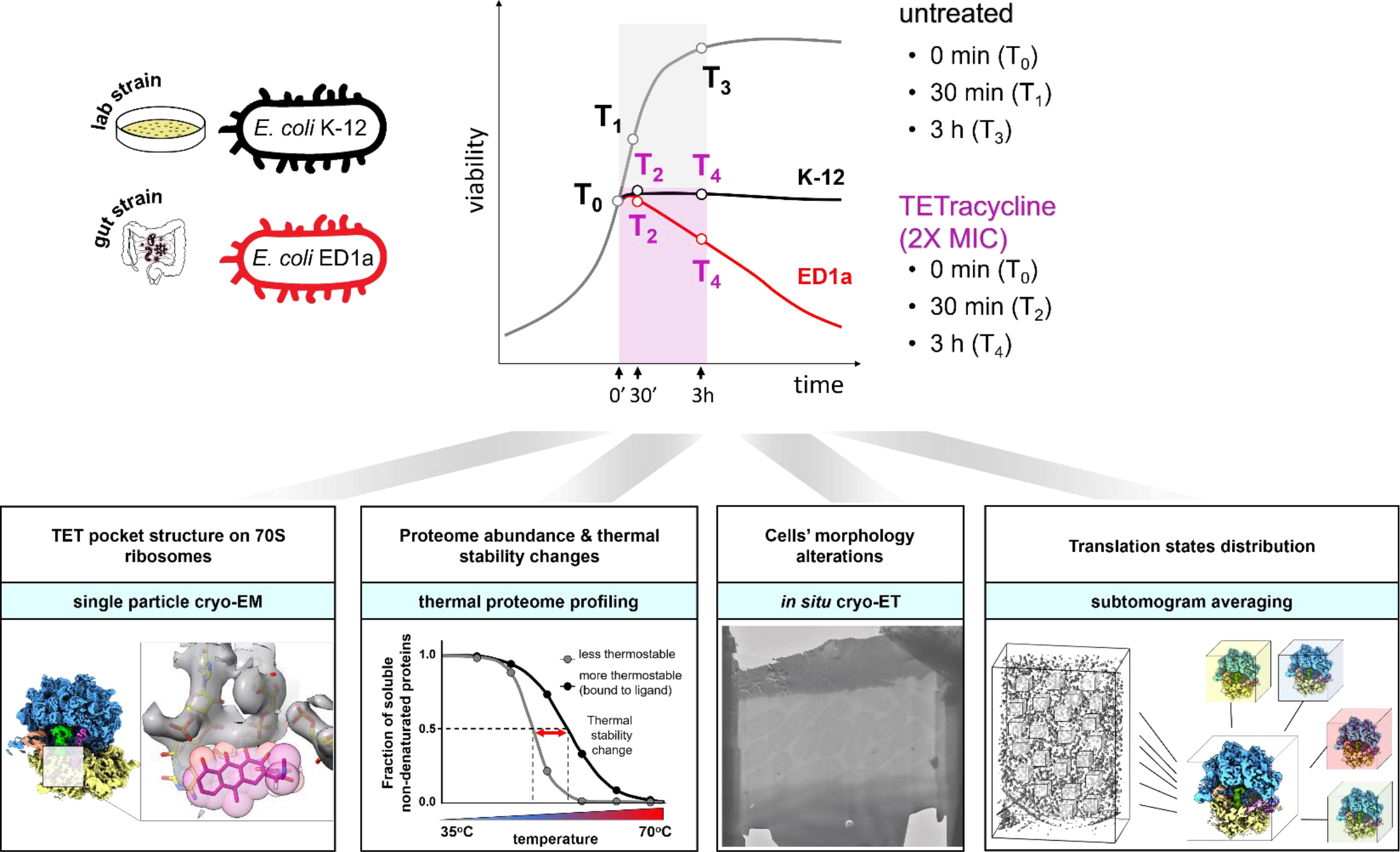
Experimental design to study bacterial response to TET. Integrative approach to investigate the short and prolonged response to TET of the lab strain *E. coli* K-12 and the gut isolate *E. coli* ED1a. Both strains were exposed to TET for 30 min and 180 min. Upper panel: a schematic viability diagram (upper right) displaying the differences in survival between two *E. coli* strains. The black line illustrates the bacteriostatic effect of TET on K-12, the red line – bactericidal effect on ED1a. The gray line illustrates the growth of untreated cells. White dots represent the timepoints used in the study. Lower panel: the structure of the respective TET binding site on the 70S ribosomes was addressed by single particle cryo-EM; changes in proteome abundance and thermal stability by thermal proteome profiling, cell morphology by in situ cryo-ET, and the translation states of ribosomes by subtomogram averaging.

## RESULTS

### TET is bactericidal for the gut isolate *E. coli* ED1a

We aimed to resolve key molecular characteristics of the antibiotic stress response differentiating two *E. coli* strains, the laboratory strain *E. coli* BW25113 (K-12), and the gut isolate *E. coli* ED1a. The experimental setup is described in a schematic illustration in **Figure 1**. A previous study reported that these two strains exhibit differential responses to the tetracycline derivative doxycycline (Maier et al., 2021). For both strains, we observed a similar effect of conventional TET at double minimal inhibitory concentration (2X MIC, which equals to 8 µg/ml for both strains). The number of culturable cells of *E. coli* K-12 did not change subsequent to exposure to TET, suggesting its bacteriostatic effect on the lab strain, while the viability of ED1a cells dropped by almost 99% after 3 hours, demonstrating a bactericidal effect of the antibiotic on the gut isolate (**Figure 2A-C**). For both strains, we measured the intracellular concentration of TET by spectrophotometry, taking advantage of a high absorbance peak of TET at 365 nm, which is otherwise absent in untreated *E. coli* lysates (**Figure S1A**). At any condition ranging from 0 to 64 µg/ml of TET added to the medium, the intracellular concentration of antibiotic was about twice as high in ED1a cells compared to K-12 (**Figure 2D**). However, even at high TET concentrations up to 4X and 8X MIC, the K-12 strain displayed a significantly higher survival rate than the ED1a strain (**Figure 2E,F**). Since even comparable intracellular TET concentrations did not lead to similarly reduced K-12 survival, these results suggest that increased accumulation of antibiotic inside the cells is not the cause for the reduced ED1a survival.

**Figure 2.**
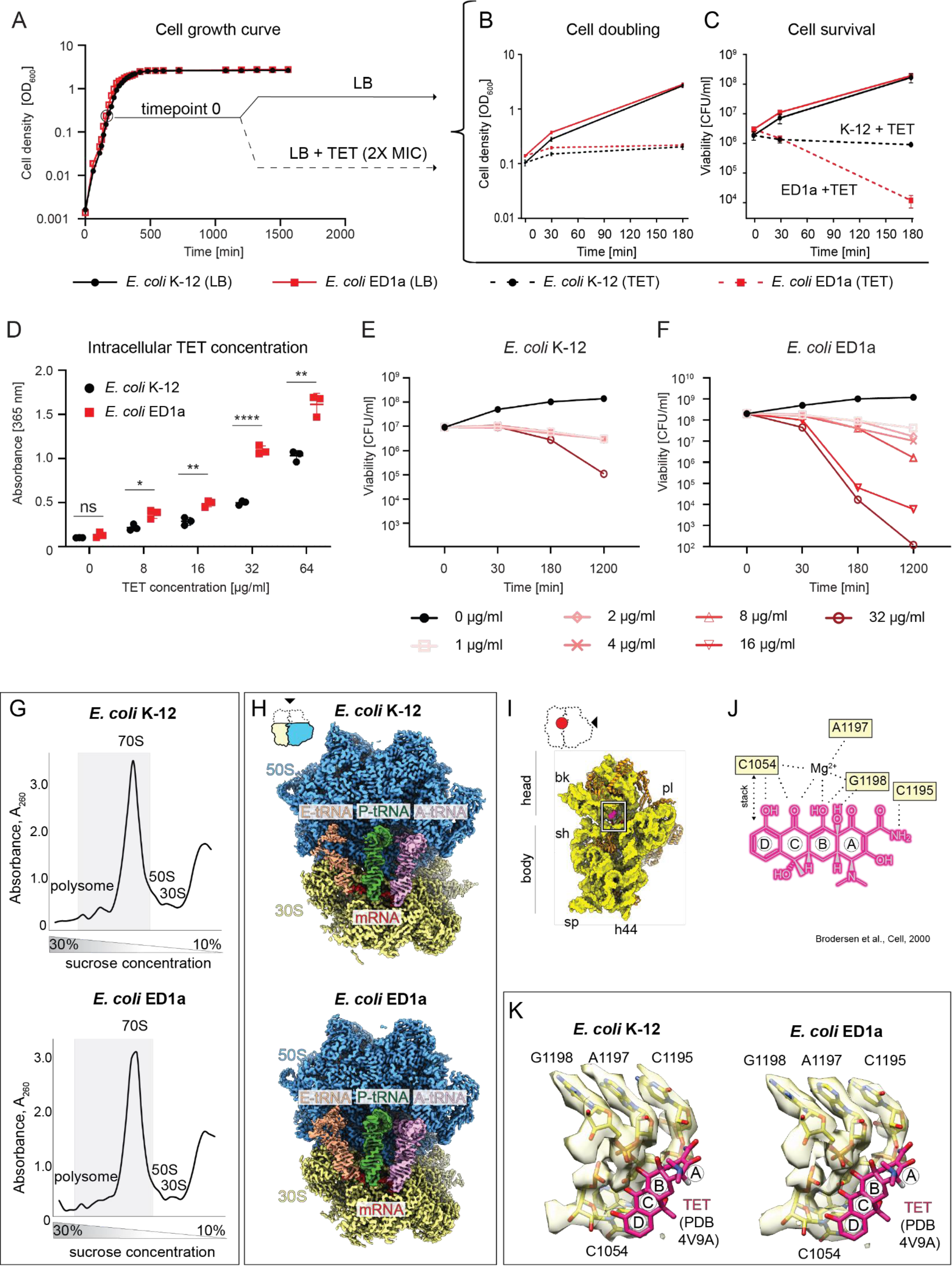
Bactericidal effect is not linked to changes of TET intracellular concentration or ribosome binding pocket. (A) Growth curves of two *E. coli* strains K-12 (black) and *E. coli* ED1a (red). The profiles show almost identical growth of the two strains. Timepoint 0 indicates the cells at exponential growth phase chosen for treatment with antibiotic in subsequent experiments. (B) The ability of cells to double in presence of TET assessed through the optical density measurements of cultures in exponential growth phase in LB (continuous lines) or in LB containing 8 μg/ml, (2X MIC) TET (dashed lines). The cells of both strains were unable to double in presence of TET. (C) Viability of cells treated with 8 μg/ml, (2X MIC) TET as in (B) assessed by counting colony-forming units (CFU) on LB (plates). The values and error bars represent averages ± two standard deviations for three independent biological replicates for each condition (of three independent measurements at every timepoint of every sample). (D) Intracellular TET concentration of cultures exposed to different TET concentrations (0 – 64 μg/ml) for 30 min, measured as absorbance of cell lysates at 365 nm. The concentration in *E. coli* K-12 (black) is almost two times lower than the *E. coli* ED1a (red). Statistical significance was calculated using a two-sided unpaired *t*-test of measurements obtained from 3 independent biological replicates. The significance symbols indicate P values: * for P ≤ 0.05, ** for P ≤ 0.005, **** for P ≤ 0.0001. (E) – (F) Viability of the *E. coli* K-12 (E) and ED1a (F) cells exposed to different TET concentrations (0 – 32 μg/ml) for up to 20 h, assessed by counting colony forming units (CFU) on LB (plates/agar). The survival of *E. coli* ED1a cells is lower than that of *E. coli* K-12. The values and error bars represent averages ± two standard deviations of three biological replicates. (G) Sucrose density gradient profiles of the clarified lysates of K-12 (top) ED1a (bottom) strains. The gray rectangle marks the fractions used for cryo-EM data collection. (H) Single particle reconstruction of 70S ribosomes from the two strains. The maps represent similar classes containing mRNA (red) and A-, P-, and E-tRNAs (pink, green, and beige respectively). (I) The subunit interface view of the 30S for *E. coli* ED1a ribosome (yellow) with TET molecule (magenta) modeled into its binding region. The 30S model of the *E. coli* K-12 70S bound to TET (PBD 5J5B) was rigid-body fitted into the single particle cryo-EM density of 70S from *E. coli* ED1a obtained in this study. The alignment was focused on the TET-binding region (nucleotides C1054, G1195, A1197, G1198). The 50S part of the model is omitted from the figure for clarity. The 16S rRNA is shown as yellow surface, the 30S r-proteins are shown as orange ribbons, the 30S parts are labeled as bk – beak, pl – platform, sh – shoulder, sp – spur, h44 – 16S rRNA helix 44. (J) Simplified scheme of TET interaction with the 16S rRNA nucleotides, adapted from (Brodersen et al., 2000). (K) The TET binding pocket of the two *E. coli* strains resolved in respective SPA structures. The 16S rRNA is colored yellow. TET molecule coordinates (magenta) were retrieved from PDB ID 4V9A and modelled in the corresponding position based on the alignment of the 16S rRNAs.

### The known TET-binding pockets are indistinguishable in K-12 and ED1a ribosomes

To test if TET could similarly bind to its target in both strains, we solved the high-resolution single-particle cryo-EM structures of the respective ribosomes and compared their TET-binding pockets. In order to obtain a close-to-native sample, we bypassed strict purification and analyzed the crude ribosome mixture pooled from the sucrose gradient sedimentation (**Figure 2G**). This allowed us to classify different ribosomal states, and we selected the natively formed 70S particles containing an A-tRNA, as the engagement of the A-site with the tRNA conveys structural stability to the TET-binding region. We resolved the structures of 70S containing mRNA and three tRNAs at 3.0 Å resolution from both strains (**Figure 2H, Figure S1B**) and found that the TET-binding pocket formed by nucleotides of the 16S rRNA of the small subunit (**Figure 2I,J**) had identical arrangements in both strains (**Figure 2K**). This region is also identical to the crystal structures of 70S-TET complexes from other bacteria (**Figure S1C**). These results strongly support the notion that the binding mode of TET to the major site at the ribosome is the same in both strains and highly conserved across bacteria.

### TPP identifies ribosomal proteins and ribosome biogenesis factors as differentially affected

To investigate further why the two strains respond differently to TET, we applied the 2D-TPP approach (Becher et al., 2018) (**Figure S2A**). This method quantifies changes in protein abundance, thermal stability, and solubility, thereby creating a holistic overview of the proteome state and revealing potential off-target effects (Savitski et al., 2014). Protein stability is a good proxy for the binding capacities of proteins for small molecules that stabilize the respective binding pockets. For both strains, we collected TPP data after 30 min or 3 h exposure to TET (8 µg/ml, 2X MIC) and from untreated cells. The proteomic screen quantified more than 1800 proteins, 1219 of which overlapped in both strains, while 318 were *E. coli* K-12-specific, and 332 were annotated as *E. coli* ED1a-specific (**Table S1, Figure S2B)**. Overall, after 30 min of TET exposure, changes in protein thermal stability were more pronounced than changes in abundance (**Figure 3A** block T2, **Figure S2C** block T2), while prolonged TET treatment resulted in more prominent changes in protein abundance (**Figure 3A** block T4, **Figure S2C** block T4). Thus, the earlier timepoint likely reflects a more immediate modulation of the thermal stability of available proteins, while cells reshape the proteome in the longer term, particularly untreated cells after 3 h, as they undergo a transition to the stationary phase (**Figure S2C** block T3). Thus, we focused our comparative analysis mostly on the short-term response (30 min), while timepoint 3 was excluded from a further comparative analysis.

**Figure 3.**
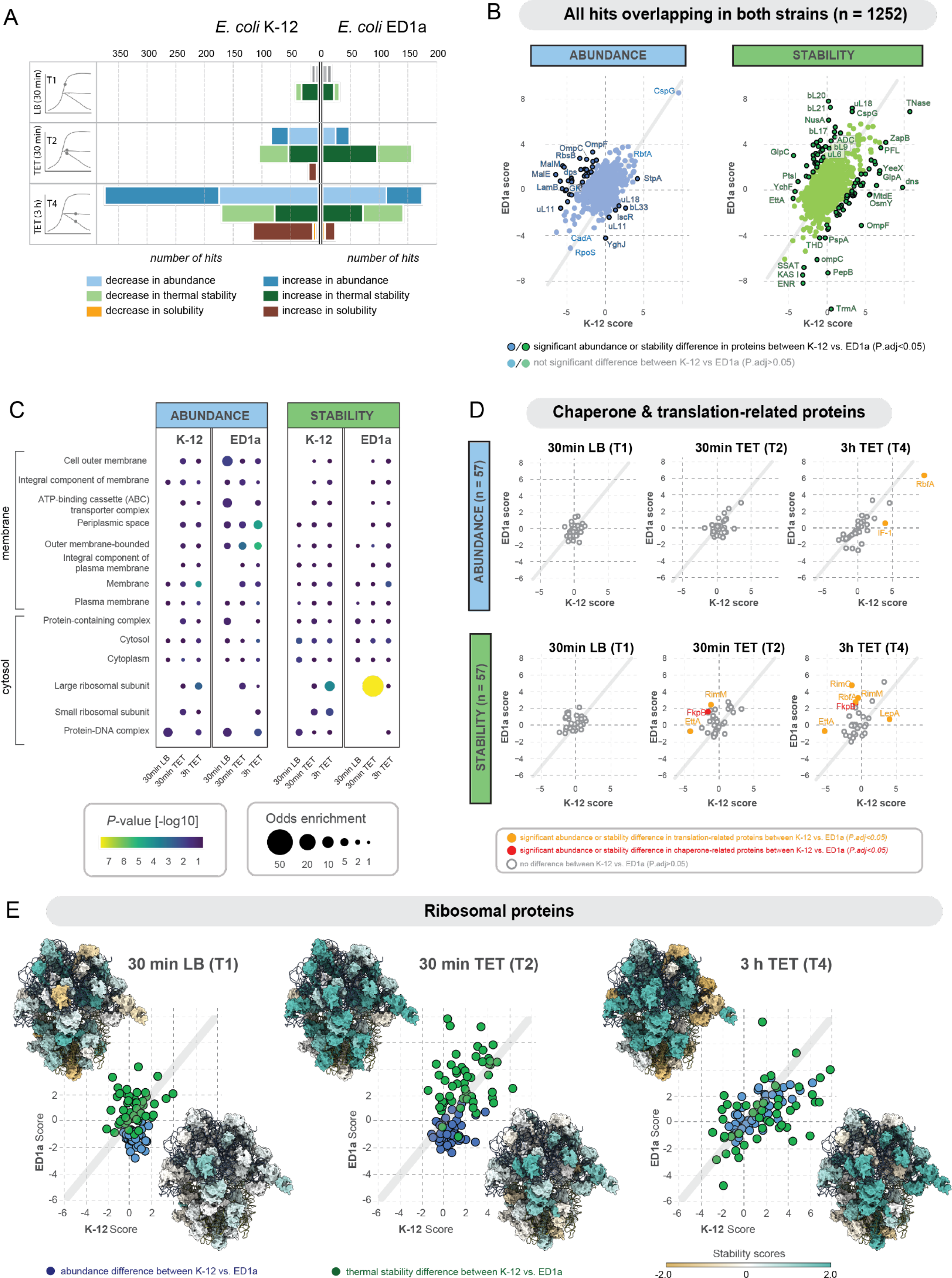
2D-TPP identifies ribosomal proteins and ribosome biogenesis factors as differently affected by TET in the two strains. (A) Comparison of the number of significantly affected proteins in abundance (blue shades), thermal stability (green shades), and solubility (orange/brown shades) for *E. coli* K-12 (left side) and *E. coli* ED1a (right side). Timepoint T1 represents untreated cells grown for 30 min, timepoints T2 and T4 represent TET-treated cells grown for 30 min and 3 h, respectively. The scores are calculated relative to the control condition T0 (untreated cells, 0 min LB). Timepoint T3 (3 h, untreated control) is shown in Fig S2C since the data were not comparable as cells shifted to stationary phase only in this set-up. (B) Scatter plot of abundance (left) and stability values (right) for TTP hits overlapping in both strains for T2 (30 min TET), comparing K-12 scores (*x*-axis) against ED1a scores (*y*-axis). Each dot represents a protein; if it is highlighted with a black contour, it indicates that there is a significant difference between K-12 and ED1a scores (*P_adj_*<0.05). The color scheme is explained in the legend below the figure panel. (C) Gene ontology (GO) enrichment plot of TPP results for different cellular compartments, with compartments ordered according to their physical location on the outer membrane – cytosol axis. The enrichment for each GO-term in the significant protein set (global & local FDR < 0.05) in each respective condition (30 min LB, 30 min TET, 3h TET) is calculated using the Fisher Exact Test, relative to the insignificant portion of proteins in each condition (displayed as color gradient for p from highest (yellow) to lowest (blue) significance). If the GO-term is enriched in at least one condition (*P*<0.01) and has at least 10 significant protein components significantly affected in the dataset, it is shown on the *y*-axis of the bubble plot. The bubble size indicates the odds enrichment, whereas the bubble color reflects the *P*-value (-log10 scale). (D) Scatter plot of abundance and stability values for proteins involved in translation and chaperoning functions for T1 (30 min LB), T2 (30 min TET), and T4 (3 h TET) respectively, comparing K-12 scores (*x*-axis) against ED1a scores (*y*-axis). Each dot represents a protein; orange and red coloring indicates a significant difference between K-12 and ED1a scores (*P_adj_*<0.05). (E) Scatter plot of abundance (blue) and stability values (green) for ribosomal proteins for T1 (30 min LB), T2 (30 min TET), and T4 (3 h TET) comparing K-12 scores (*x*-axis) against ED1a scores (*y*-axis). Each dot represents a protein. The two insets in each plot depict 70S ribosome structures shaded according to the respective thermal stability score values of the respective protein components (upper left corner: coloring based on ED1a scores; lower right corner: coloring based on K-12 scores). The PDB model was combined from coordinates fetched from PDB 3J7Z, PDB 7K00, and PDB 6H4N. Proteins with increased stability are colored turquoise, proteins with decreased stability are colored tan, and proteins with unchanged stability are colored white.

Both strains showed a significant increase in abundance of cold-shock proteins, including RbfA, CspB, and CspG (**Figure 3C**). Earlier, similar effects were reported for TET-treated *E. coli* K-12 by 2D electrophoresis (VanBogelen and Neidhardt, 1990) indicating that our approach and results are in line with prior biochemical studies. At the same time, we detect notable differences between the strains in abundance or stability scores for a number of proteins. Among them, the proteins of malate metabolism (MalE, MalM), and galactose metabolism (GatA, GatC, gatD, GatZ), and several translation-associated proteins including EttA, TrmA, RimO, translation initiation factor IF-1 (**Figure 3B)**. Noticeably, we detect strong changes in abundance or thermal stability over the course of TET treatment (T2 vs T4) for several proteins annotated only in one of the strains, including several uncharacterized proteins **(Figure S2D-E)**. We designate these hits as strain-specific proteins.

Gene ontology term analysis identified ribosomal proteins (r-proteins), in particular components of the 50S large ribosomal subunit, as the major category affected by TET in terms of thermal stability (**Table 1**, **Figure 3C, Figure S2F**). Additionally, ribosome biogenesis factors, such as RimM, RimO, EttA displayed a significant increase of thermal stability scores in TET-treated ED1a cells (**Figure 3D)**. Notably, these, as well as r-proteins themselves, only showed high thermal stability fluctuations, whereas the abundance remained mostly unchanged over time and regardless of antibiotic treatment (**Figure 3D,E, S3A-C**). This suggests that the number of ribosomes remained constant in all conditions, but that their structural status was altered shortly after treatment, specifically for *E. coli* ED1a (**Figure 3E, S3B**). Thermal stability scores increased particularly for proteins near the subunit interface, an effect that may be attributed to reduced inter-subunit rotation (**Figure S3B**). Overall, these data identify stress response, metabolic and translation factors to be affected by treatment, and underscore a stronger impact of the drug in ED1a cells.

### Cryo-ET visualizes distortion of the bacterial envelope in the gut strain

Recent studies demonstrated the power of subtomogram averaging to determine high-resolution structures of ribosomes and to resolve their different conformations *in situ* (Hoffmann et al., 2022; Xing et al., 2023; Xue et al., 2022). We performed *in situ* cryo-ET coupled to subtomogram averaging (STA) to visualize cell morphology and the structural state of the protein synthesis apparatus in the two *E. coli* strains upon TET treatment. We obtained lamellae using the focused ion beam (FIB) milling technique and collected the tilt series of individual bacterial cells at 30 min without treatment (30 min LB), and treated (30 min TET) (**Figure 4A-C**).

**Figure 4.**
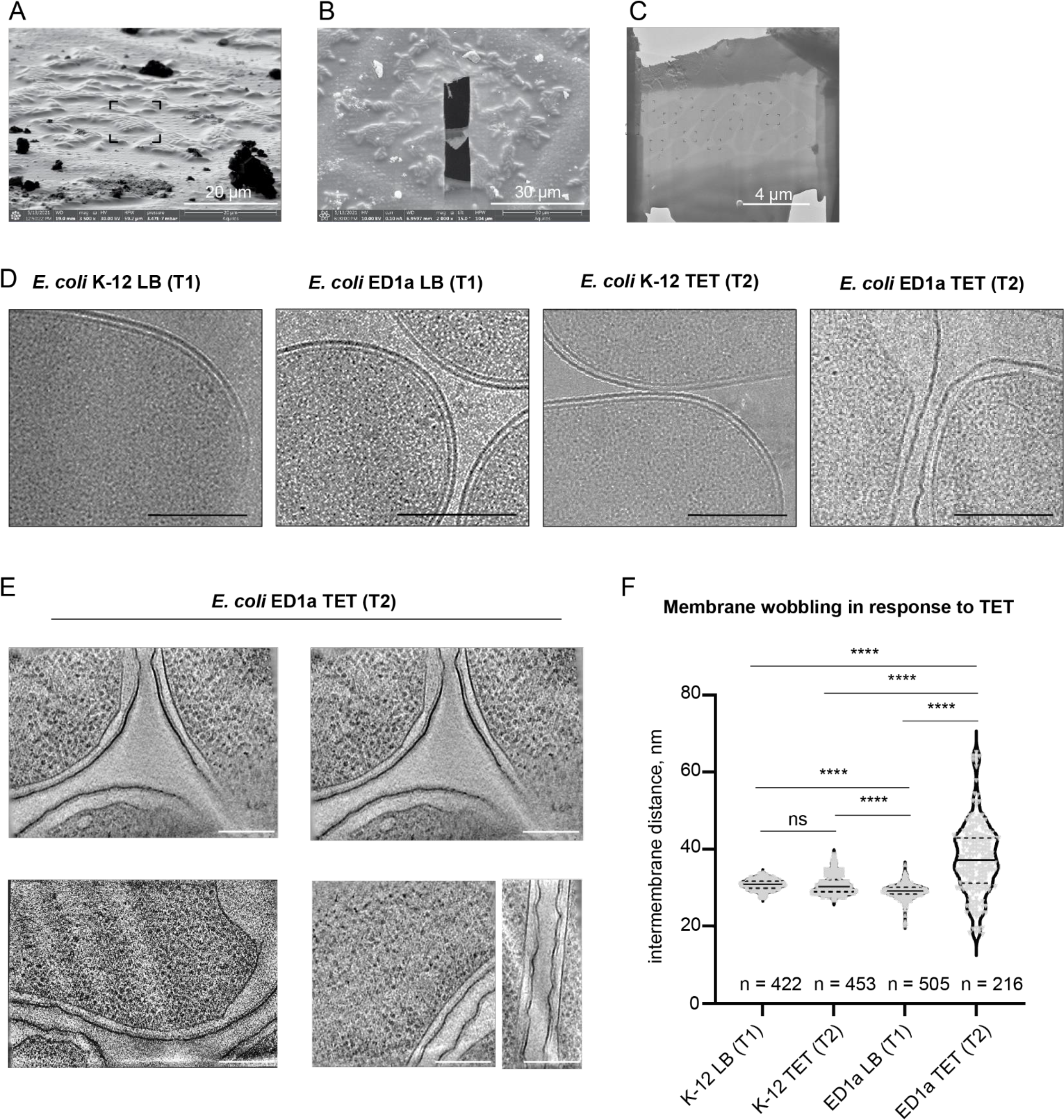
Cryo-ET reveals severe membrane wobbling in *E. coli* ED1a cells treated with TET. (A) Ion beam image of the *E. coli* cells selected for FIB-milling. Scale bar 20 µm. (B) Scanning electron beam image of the grid square with FIB-milled lamella of bacterial cells. Scale bar 30µm. (C) Transmission electron microscopy (TEM) image map of a lamella. The black targets point to the regions of subsequent tilt series data collection. Scale bar 4 µm. (D) Snapshots from the TEM lamellae maps acquired at 6,500 X magnification, demonstrating the membranes wobbling in *E. coli* ED1a cells treated with 8 μg/ml (2X MIC) TET for 30 min. Scale bar 0.5 μm. T1: timepoint 1 (30 min LB), T2: timepoint 2 (30 min TET). (E) Digital slices at different z-heights from tomograms of *E. coli* ED1a cells treated with TET for 30 min. The images were low pass filtered to better visualize the membrane wobbling. Scale bars 0.2 μm. (F) Intermembrane distance in the cells used for cryo-ET and subtomogram averaging. The calculations were performed on the visible part of the cell envelope extracted directly from the tilt series collected at 53,000 X magnification. The distances between two membrane leaflets were measured at points spaced by 10 nm using a script (https://github.com/martinschorb/membranedist) and manually inspected to remove obvious outliers. Intermembrane distances were higher on average and significantly more variable in the *E. coli* ED1a cells treated with TET for 30 min. The violin plot shows the smoothed density of every measurement (black dots), the first and third quartiles (black solid lines), and the median (black dashed line); *n* represents the number of points at which measurements were made for 10 K-12 (LB), 10 K-12 (TET), 10 ED1a (LB), and 7 ED1a (TET) cells. The violin plots for each of these cells individually is shown in Figure S4A-D. The schematic cartoon of intermembrane distance analysis is shown in Figure S4E. The statistical significance was calculated using a two-sided unpaired *t*-test. **** signifies a *P*-value<0.001.

Visual inspection revealed that TET-treated *E. coli* ED1a cells had a distorted cell envelope, in particular wobbling of outer and inner membranes (**Figure 4D,E**). We measured the periplasmic thickness of the cells using computational segmentation of membranes and calculated the distance between inner and outer membranes as previously described (Zietek et al., 2022). Upon treatment with TET, the uniformity of periplasmic space of *E. coli* ED1a cells was significantly perturbed, which resulted in overall longer intermembrane distances across the cells and irregularity within individual cells (**Figure 4F; Figure S4A-D**). Although the colony counting assay showed that at least half of ED1a population was still viable after 30 min of TET treatment (**Figure 2C**), the envelop disruptions were visualized in virtually every cell analysed by cryo-ET, suggesting that the damage of some cells was still below a threshold allowing recovery. This is in accord with the TPP data that had identified porins OmpF, OmpC; multidrug efflux pumps AcrAB-TolC, MdtE, EmrA; ABC (ATP-binding cassette) transporters Dpp, Opp, MsbA, Mal, Rbs; and cell envelope-shaping proteins, OmpA, TolR to differ in abundance and thermal stability between both strains (**Table S1, Figure S2D**). Based on similar observations, TET or its analogs were previously suggested to have an additional mode of action, independent of translation inhibition (Oliva et al., 1992; Wenzel et al., 2021).

### The majority of ribosomes in the gut strain are in a translation incompetent state upon TET treatment

To obtain spatial and temporal information on the 70S ribosomes in untreated and treated cells, we applied the STA approach. Considering the high degree of structural similarity observed in our cryo-EM structures of ribosomes in the two strains, we merged 70S subtomograms from all datasets to obtain a resolution benchmark for our STA pipeline (**Figure 5A,B, Figure S5**). The resulting subtomogram average of *E. coli* 70S ribosomes overall reached a resolution of 5.2 Å (**Figure 5C, Figure S5**). Some regions of the structure were resolved to side-chain details. Among them, Lys44 and Arg46 of the protein uS4 that supports placement of mRNA at the mRNA entry pore formed by uS3, uS4, and uS5, were defined at this depth (**Figure 5D**).

**Figure 5.**
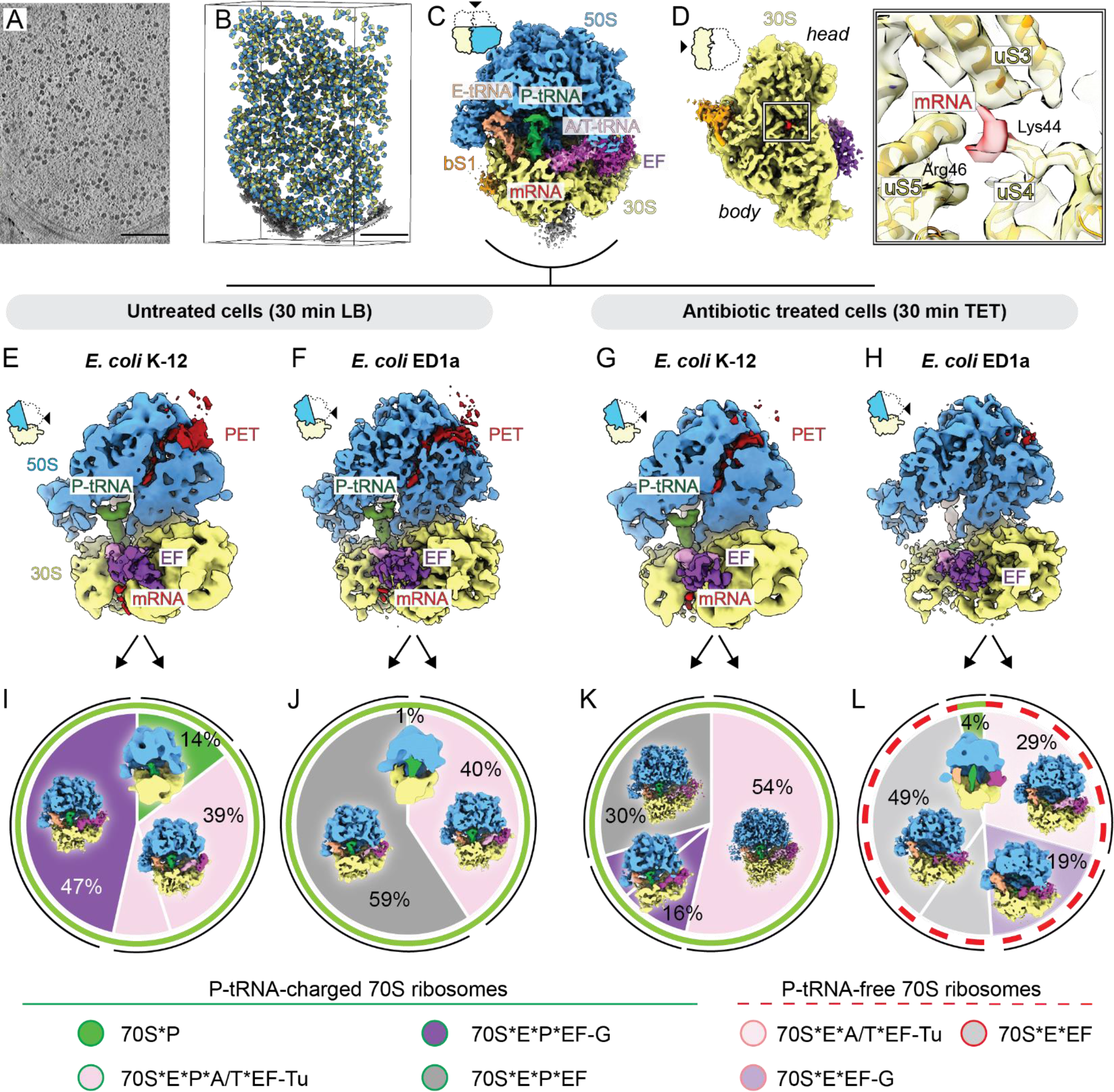
In situ cryo-ET, subtomogram averaging and 3D classification reveals majority of *E.coli* ED1a 70S ribosomes are in translation-incompetent state lacking P-tRNA after TET treatment. (A) Digital slice from a representative tomogram of *E. coli* ED1a cells. The tomogram was denoised and the contrast was enhanced using IsoNet (Liu et al., 2022) trained on 5 tomograms with different defocus. Ribosomes appear as dark spherical objects. Scale bars 0.2 μm. (B) Three-dimensional view of the tomogram from (A) with aligned ribosome particles projected back into tomographic volumes. Small ribosomal subunits colored yellow, large subunits colored blue. Membranes segmented from tomogram are colored in gray. (C) The subtomogram average of the *E. coli* 70S ribosome from all datasets was resolved at 5.3 Å resolution. The structure represents an actively translating ribosome with mRNA (red), A-, P-, E-tRNA ligands, (pink, green, and beige respectively), elongation factors (EF, magenta) bound. (D) The solvent side view of the small ribosomal subunit of the overall subtomogram average of *E. coli* 70S ribosome. The enlarged inset shows details of the mRNA entry tunnel composed of universal r-proteins uS3, uS4, and uS5. The mRNA is supported by Lys44, and Arg46 of uS4, for which density is resolved to side-chain resolution. The segmented density for r-proteins shown in yellow, for mRNA in red. For visualization reasons, and to make the mRNA entry tunnel visible, the extra density at the polysome interface between the head and the body of the 30S is omitted. It can be seen colored in gray in figure panel C. (E) – (H) Subtomogram averages of the 70S ribosome from (E) – *E. coli* K-12 (LB) at 8.5 Å, (F) – *E. coli* ED1a (LB) at 7.5 Å, (G) – *E. coli* K-12 (TET) at 8.5 Å, (H) – *E. coli* ED1a (TET) at 7.0 Å. The structure from the *E. coli* ED1a TET sample (H) shows only negligible density for tRNA and peptide exit tunnel (PET), suggesting the translation-incompetent state of most of the ribosomes in these cells. The 50S subunit is clipped to visualize the nascent chain inside PET (dark red), and the P-tRNA (green). Ribosomal subunits and ligands coloring: 30S – yellow, 50S – blue, mRNA – red, A-, P-, E-tRNA ligands – pink, green, and beige respectively, elongation factors (EF) – magenta. (I) – (L) The distribution of ribosome classes sampled from *E. coli* K-12 30 min LB (I), *E. coli* K-12 30 min TET (J), *E. coli* ED1a 30 min LB (K), *E. coli* ED1a 30 min TET (L). Each segment of the pie chart represents a separate class obtained by 3D classification: 70S*P complex – green, 70S*E*P*A/T*EF-Tu complex – pink, 70S*E*P*EF-G complex – purple, 70S*E*P complex with weakly resolved elongation factor (EF) – gray, 70S*E*A/T*EF-Tu complex – light pink, 70S*E*EF-G complex – orchid, 70S*E complex with weakly resolved elongation factor (EF) – light gray. The thick green contour defines ribosome classes of translation-competent ribosomes charged with P-tRNA, the thick red dashed contour defines classes of translation incompetent ribosomes lacking P-tRNA. The thin black contours outline structurally similar 3D classes.

We then determined 70S ribosome structures from *E. coli* K-12 and *E. coli* ED1a for control (30 min LB) and 30 min TET-treated cells individually at a resolution range from 7.0 to 8.5 Å (**Figure 5E-H**). To disentangle the different functional states of ribosomes in each of the four conditions, we performed multiple rounds of 3D classification and sampled ribosome structure heterogeneity based on ligand occupancy (**Figure S6A-D**). Without the antibiotic, we observed the densities attributed to mRNA, P-site tRNA and the nascent peptide similarly for both strains (**Figure 5E,F**). The major classes in untreated samples represented 70S·E·P·A/T-tRNA·EF-Tu complex and 70S·E·P-tRNA·EF-G complex (**Figure S6A,B**), resembling the slow translation states of tRNA accommodation and peptide bond formation (Gromadski and Rodnina, 2004; Rodnina, 2018; Savelsbergh et al., 2003). For the K-12 30 min TET sample, we observed similar ligand composition (**Figure 4G**), and all classes contained P-tRNA and elongation factors, suggesting that these ribosomes are translation competent (**Figure S6C**). In striking contrast, the densities for the mRNA, P-tRNA, and the nascent chain were absent in 70S of *E. coli* ED1a 30 min TET (**Figure 5H**), implying differences in the translation capacity. Subsequently, 3D classification revealed that 96% of TET-treated ED1a ribosomes do not carry P-tRNA, a ribosomal state that is incompatible with protein synthesis. A large proportion of ribosomes in this sample were annotated as 70S·A/T·E-tRNA·EF-Tu complexes (**Figure S6D**). Canonically, a complex of EF-Tu with aminoacyl-tRNA (aa-tRNA) binds to the ribosome to promote the elongation of the nascent chain bound to the P-tRNA (Voorhees and Ramakrishnan, 2013). The observed presence of EF-Tu on the P-tRNA-deficient 70S could potentially be caused by the generation of run-off ribosomes that could not be recycled for translation initiation.

Surprisingly, regardless of TET treatment 70S averages contained density for the ribosomal protein bS1, with the resolved N-terminal ribosome anchor and the domain I regions (**Figure S6E**). This suggests that after aiding mRNA accommodation on the 30S subunit (Duval et al., 2013; Qu et al., 2012), bS1 remained bound to the ribosome. This finding corroborates the vital role of this protein in *E. coli* translation (Sorensen et al., 1998) and indicates that binding may be required for the later stages of the translation cycle, transcription-translation coupling (Demo et al., 2017), or ribosome hibernation (Beckert et al., 2018). Overall, these results suggest that after 30 min TET treatment *E. coli* ED1a contains a very small pool of ribosomes that are potentially capable of protein synthesis. This is in accord with the results of the TPP screen, which show an increase of thermal stability of ribosomal proteins at the subunit interface, thus indicating that ribosomes are present in a structurally inflexible state.

## DISCUSSION

Tetracyclines are by and large considered bacteriostatic antibiotics, yet the two *E. coli* strains analyzed in our study exhibit differential responses, with bactericidal activity seen in ED1a (**Figure 2C,F**) (Maier et al., 2021). It has previously been postulated that ribosome-binding antibiotics can be bactericidal for cells with few ribosomal operons, which also have reduced growth rates (Levin et al., 2017). However, considering that both *E. coli* K-12 and *E. coli* ED1a have seven ribosomal operons and a comparable growth rate (**Figure 2A**), we concluded that the killing of the gut isolate by TET is unlikely to be linked to altered rRNA transcription. Moreover, the number of ribosomes per cell detected by cryo-ET is similar in both strains (**Figure S5**). The observed increased intracellular antibiotic concentration in *E. coli* ED1a (**Figure 2D**) does not appear to be the main causative effect, as treatment of *E. coli* K-12 with 16 ug/ml TET, a concentration which results in a similar intracellular concentration to ED1a treated with 8 ug/ml (**Figure 2D-F**), retains the bacteriostatic phenotype. Direct binding of TET to the 30S A-site of the bacterial ribosome is well documented (Cocozaki et al., 2016; Jenner et al., 2013), however, we find that this canonical binding site alone is unlikely the singular determinant for susceptibility towards treatment, as single-particle cryo-EM structures of 70S show entirely indistinguishable TET binding sites in both strains (**Figure 2K**). Together these findings demonstrate that the effect of TET on these bacterial cells is more complex and that multiple aspects likely have to be considered to explain the overall phenotypic differences observed.

In addition to pointing towards an involvement of ribosomal biogenesis, translation, and protein folding (**Figure 3C-E, Figure S3A-C**), we detected subtle differences in the proteomic makeup of membrane proteins, including proteins that have been previously associated with antibiotic response (**Table S1, Figure S2D**). Previous proteomic and mutagenesis/genetic analyses of *E. coli* K-12, *Acinetobacter baumannii* DU202, and an environmental-borne *E. coli* EcAmb278 also found an alteration in the abundance of several membrane transporters, including OmpC, TolC, LamB and peptidoglycan-based cell wall proteins upon TET treatment (Jones-Dias et al., 2017; Xu et al., 2006; Yun et al., 2008; Zhang et al., 2008). The differential effects we detected in the two strains by proteomics could be related to the phenotypic differences observed in membrane morphology by cryo-ET (**Figure 4D,E**). In our large-scale 2D-TPP screen, we identify numerous species-specific proteins and putative uncharacterized proteins with significant changes in their abundance and thermal stability upon TET treatment (**Table S1**, **Figure S2E**). These include several proteins with yet unknown or poorly described function, where a potential role in the response to TET would need to be further assessed.

The ability of cells to cope with stress conditions and to quickly recover, strongly depends on the activity of the translation machinery, and the functional state of the ribosomes (Cheng-Guang and Gualerzi, 2021). Using cryo-ET, we find that in contrast to *E. coli* K-12, in *E. coli* ED1a cells, TET treatment leads to presence of almost all ribosomes (96%) in a very specific state of 70S containing translation factors and E-tRNA, but lacking the P-tRNA (**Figure 5L, Figure S6E**). This conformational state is incompatible with translation, as the P-tRNA retains/locks the nascent peptide chain on the ribosome and is required for every round of incorporation of new amino acid. In the recent in situ cryo-ET studies of *Mycoplasma pneumonia* bacterium treated with other ribosome-binding antibiotics such as chloramphenicol or spectinomycin, the P-tRNA-lacking state of the ribosome was not observed (Tegunov et al., 2021; Xue et al., 2022). A similar approach in untreated eukaryotic cells found a fraction of ribosomes with the vacant P-tRNA site, but determined these as being in a hibernating-like state (Hoffmann et al., 2022; Xing et al., 2023). Moreover, in human cells treated with translation inhibitor homoharringtonine, only 47 % of ribosomes turned into P-tRNA-deficient state, despite showing the drug bound to the majority of all ribosome particles (Xing et al., 2023). All this suggests that the response of *E. coli* ED1a to TET may be more complex and not depend only on translation regulation, but also other factors which have yet to be discovered. Considering the recovery rate of only about 50% of ED1a cell population after 30 min, and almost complete death after 3 h of TET treatment (**Figure 2C**), the observed translation-incompetent state might be irreversible. In contrast, P-tRNA is present in every class in the K-12 TET sample, indicating that these ribosomes likely remain translation competent, and possibly restart protein synthesis upon TET removal. This could then be very rapid, as our cryo-ET structures show that in *E. coli* K-12 cells, the EF-Tu·aa-tRNA complex can bind to the 70S ribosome even in presence of TET (**Figure 4E)**, suggesting a high competition between reversible antibiotic and the tRNA binding to the ribosomal A-site inside cells. All this could potentially provide a high survival to the *E. coli* K-12 strain under TET pressure.

Lastly, in our cryo-ET data, we did not observe any class of protein-induced inactive ribosomes, such as 50S·RsfS complexes (Khusainov et al., 2020) or 100S and 70S complexes (Ortiz et al., 2007; Polikanov et al., 2012), which are mediated by hibernation promoting factor HPF. We also did not detect significant changes in abundance or thermal stability of HPF protein by proteomics (**Table S1**). Although the HPF binding site is known to overlap with that of TET (Khusainov et al., 2017) and thus could protect ribosomes by reversible binding, our results indicate that ribosomes are preferentially retained in the pro-active 70S form upon TET treatment. This is true even in the ED1a strain, which experiences a dramatic conversion of nearly all ribosomes to a translation-incompetent state, which is potentially not reversible.

From a technical perspective, most of bacterial species, including *E. coli*, are too thick for the direct imaging by transmission electron microscopy, and require thinning by focused ion beam, similar to any eukaryotic cell. Moreover, the dense molecular crowding in the cytosol and the absence of cellular compartmentalization make *E. coli* a challenging sample for cryo-ET tilt-series alignment and high-resolution subtomogram averaging. Such limitations likely restricted many cryo-ET studies on bacteria to the analysis of phenotypic features, or structure determination of peripheral macromolecular complexes (reviewed in (Khanna and Villa, 2022)). On top of that, quantitative structural analysis of the highly dynamic molecular machineries such as ribosome require a very delicate sample preparation with no sample intervention before plunge-freezing. In this study, we established a protocol for close-to-native *E. coli* sample preparation for cryo-ET (Methods) suitable for high-resolution subtomogram averaging of cytosolic complexes and thereby provide the first high-resolution *in situ* snapshots of ribosomes and the structural malfunctions induced by TET both in lab *E. coli* strains and in human gut isolate.

Our study shows that *in situ* and in-depth molecular analyses are essential to further advance the overall understanding of antibiotic susceptibility, in particular of human commensal bacteria that may be differentially affected by antibiotic treatment. Given the increasing evidence for a vital role of the microbiota in the maintenance of human health (Gilbert et al., 2018), preserving its homeostasis and minimizing interference with its species diversity is highly important, and the development of efficient species-specific antibiotics with reduced side effects represents one of the main challenges.

## Materials & Methods

### Growth and viability assays of E. coli treated with TET at 2X MIC

For all experiments, we used the standard lab strain *E. coli* BW25113 (K-12) and a gut strain *E. coli* ED1a available in the collection of Dr. A. Typas lab (EMBL-Heidelberg). Cells were grown in 250 ml flasks containing 50 ml LB at 37 °C and 180 rpm shaking. For growth curve estimations, the cells were grown continuously for 26 hours. For the viability assays, 50 ml of cell cultures were inoculated with the overnight pre-culture to the starting OD_600_ of 0.005, and grown for 3 hours in LB until OD_600_ of 0.3, then 20 ml of these cultures were transferred to new flasks containing 20 ml of prewarmed LB or LB supplemented with 16 ug/ml tetracycline (TET). These dilutions were made to prevent the transition of control LB-grown cells to the stationary phase during the first 30 minutes. The cultures were further grown for 30 minutes and 3 h. At every timepoint, 1 ml of culture was taken for OD measurements, and 1 ml for colony forming units (CFU) estimation. To address cell viability (CFU/ml) at 8 µg/ml TET, 1 ml of cultures were washed twice in 1 ml of phosphate buffer saline (PBS) and subjected to serial dilutions in PBS. The drops of 5 ul were placed on the LB-agar plates and incubated overnight at 37 °C. To estimate viability at different TET concentrations, cell cultures of 10 mL were grown in 15 ml tubes. At every timepoint, 0.5 ml of cultures were taken for OD measurements and CFU estimation. Cells were washed twice in PBS and plated on a solid LB-agar medium as described above.

### Intracellular TET measurements

Cell cultures of 50 mL were inoculated with the overnight pre-culture to the starting OD_600_ of 0.005, and grown for 3 hours in LB until OD_600_ of 0.3. Then, 20 ml of cultures were diluted twice in prewarmed LB or LB supplemented with 16 µg/ml tetracycline (TET). After an additional 30 minutes of incubation, the cells were collected by centrifugation at 4000 × g and 4 °C for 10 min. Cells were washed twice in 2 ml of PBS, suspended in 100 ul of PBS, and lysed by heating at 100 °C for 5 minutes. To check antibiotic thermal stability, TET dissolved in PBS was also heated for 5 minutes at 100 °C and its absorbance spectra of serial dilutions were measured in NanoDrop device (Thermo Scientific). To measure intracellular antibiotic concentration in cultures treated with different initial TET concentrations (from 0 to 64 µg/ml), the 50 ml cultures grown in 250 ml flasks for 3 hours in LB were split into 5 aliquots of 10 ml and diluted twice with LB supplemented with an appropriate amount of TET. The cultures were further incubated for 30 minutes and lysed as described above.

### Ribosome purification

Both strains, *E. coli* K-12 and *E. coli* ED1a were grown for 3 h until reaching OD_600_ of 0.3 in 1 L flasks filled with 200 ml of LB. Cells were collected in 400 ml centrifuge tubes filled with 150 g of frozen buffer A (10 mM HEPES-KOH pH 7.5, 100 mM NH_4_Cl, 15 mM Mg-acetate, 1 mM ethylenediaminetetraacetic acid (EDTA), 1 mM DTT), and pelleted at 4,000 × g for 20 min at 4°. Pellets were resuspended in 7 ml of buffer A supplemented with the addition of 200 μl of protease inhibitor cocktail (one tablet (Roche), dissolved in 1 ml buffer A), of 50U DNase I (Roche) and lysed by three passages through French Press at 1000 Bar. Cell debris was removed by centrifugation at 30 000 × *g* for 60 min. The resulting supernatant was diluted to 1.5 mg/ml and 0.2 ml was layered on 10-30 % sucrose gradients prepared in buffer E (10 mM Hepes-KOH pH 7.5, 60 mM KCl, 15 mM Mg-acetate, 0.5 mM EDTA, 1 mM DTT) and spun at 32,600 × g for 15 h at 4° in SW32 rotor (Beckman Coulter). Ribosome fractions were pooled as shown in **Figure 2D**. The sucrose was removed and ribosomes were concentrated to 10 mg/ml using Amicon centrifugal filters with molecular weight cut-off 100 kDa. Aliquots of 10 μl were flash-frozen in liquid nitrogen, and stored at −80°.

### Single particle cryo-EM grids preparation and data collection

The 70S (3 μl at 1 mg/ml concentration) were applied to Quantifoil R 1.2/1.3 300 mesh grids discharged for 90 sec at 15 mA and 0.38 mbar using PELCO easiGlow. The sample was mounted on Vitrobot Mark IV (Thermo Fisher Scientific) at 100% humidity and 10 °C, then blotted for 3 s at nominal blot force 4 and plunge-frozen in liquid ethane, cooled by liquid nitrogen. Single particle cryo-EM data were collected using EPU software v2.14 (Thermo Scientific) on a 300 kV Titan Krios transmission electron microscope (ThermoFisher Scientific) equipped with a BioQuantum-K3 imaging filter at the in-house electron microscopy facility of Max-Planck Institute of Biophysics. Dose-fractionated movies were acquired in electron counting mode at 105,000 X magnification on a K3 camera, corresponding to a pixel size of 0.837 Å/pix. The calculated total electron exposure per image was 40 e^−^/Å^2^ on the specimen and the exposure rate was set to 15 e^−^/pix/sec on the camera. A total of 12,022 images were collected for *E. coli* K-12 samples, and 16,559 images for *E. coli* ED1a samples. CryoSPARC Live (Punjani et al., 2017) was used for on-the-fly data quality assessment.

### Single particle cryo-EM image processing and model fitting

Image processing was performed in CryoSPARC v3.3.1 (Punjani et al., 2017). After patch motion correction and contrast transfer function (CTF) estimation, all micrographs were kept for further analysis. A total of 1,740,048 / 2,302,674 (*E. coli* K-12 / *E. coli* ED1a) particles were picked using the blob picker with minimum and maximum particle diameters of 200 Å and 300 Å respectively, and extracted using a box size of 64 pixels at 3.348 Å/pix. The 2D classification was used to remove junk particles resulting in 1,211,457 / 1,167,533 particles that were further used for further classification. Using 70S ribosome volume generated by ab initio reconstruction from 10,000 randomly selected particles, the heterogeneous refinement was used to remove poorly aligned volumes. The remaining particles were re-extracted with the box size of 200 pixels at 1.674 Å/pix. A new ab initio model was generated, and another round of heterogeneous refinement was performed to reveal conformational heterogeneity of ribosomes. Particles of the 70S·mRNA·APE-tRNA complexes (80,169 / 132,290 particles for K-12 / ED1a respectively) were extracted with a box size of 300 pixels at 1.256 Å/pix and subjected to homogeneous refinement with default parameters. The final resolution was 3.02 Å, and 3.01 Å for K-12 and ED1a respectively. An overview of the single particle cryo-EM processing workflow is shown in Figure S2. The model of *E. coli* 70S·mRNA·APE-tRNA·paromomycin (PDB ID: 7K00) complex was rigid-body fitted into obtained experimental density maps. The 16S rRNA nucleotides around the TET-binding region were manually refined using Coot (Emsley and Cowtan, 2004). The resulting model was aligned with *E. coli* 70S·TET complex from *E. coli* (PDB ID: 5J5B) and *Thermus thermophilus* (PDB ID: 4V9A) models in ChimeraX (Pettersen et al., 2021) using MatchMaker restricted to TET-Binding region.

### Thermal protein profiling and sample preparation for MS

Thermal proteome profiling was done as previously described (Becher et al., 2018) with adaptation of cell growth conditions to our current experimental setup. Cultures were inoculated with the ov ernight pre-culture to the starting OD_600_ of 0.005, and grown for 3 hours in LB until reaching OD_600_ of 0.3. The cultures corresponding to each timepoint were grown in individual flasks. To ensure comparable total cell numbers and final amount of protein extracts in treated and untreated cultures, the cells were grown in different volumes. The total volumes of each culture were as follows (timepoint – ml): T0 – 350, T1 – 200, T2 – 500, T3 – 100, T4 – 200. For harvesting cells were transferred on ice into 400 ml centrifuge bottles, washed twice with PBS by centrifugation at 32,600 × g for 20 min at 4°, resuspended in lysis buffer (final concentration 0.8% NP-40, 1.5 mM MgCl_2_, protease inhibitor, phosphatase inhibitor, 0.4 U/μl benzonase), transferred to 2 ml tubes supplemented with glass beads. The cells were lysed by 2 cycles of 2 min vortexing at 4 °C to avoid sample heating. Cell debris was removed by centrifugation at 15,000 × g for 10 min at 4 °C. The supernatant was diluted to 1 mg/ml protein concentration and distributed by 0.1 mg in PCR tubes. Each aliquot was heated for three minutes to a different temperature (37.0-40.4-44.0-46.9-49.8-52.9-55.5-58.6-62.0-66.3°C). Protein aggregates were removed by centrifugation at 80,000 × g for 40 min at 4 °C. Protein concentration in the suparnatant was determined, and 10 μg protein (based on the two lowest temperatures) was used for further sample preparation. Proteins were reduced, alkylated, and digested with trypsin/Lys-C using the S-trap from OASIS. Peptides were labeled with TMT10plex (Thermo Fisher Scientific). Samples were combined in the following manner: experimental conditions (incl. control, T1, T2, T3, T4) were combined for each temperature resulting in five conditions analyzed together. Pooled samples were fractionated on a reversed-phase C18 system running under high pH conditions. In addition, both samples from NP-40 lysis (37°C) and SDS lysis from each experimental condition were combined, allowing the determination of protein solubility differences.

### LC-MS/MS measurement

Peptides were separated using an UltiMate 3000 RSLC nano LC system (Thermo Fisher Scientific) equipped with a trapping cartridge (Precolumn C18 PepMap 100, 5 μm, 300 μm i.d. x 5 mm, 100 Å) and an analytical column (Acclaim PepMap 100, 75 μm x 50 cm C18, 3 μm, 100 Å). The LC system was directly coupled to a Q Exactive Plus mass spectrometer (Thermo Fisher Scientific) using a Nanospray-Flex ion source. Solvent A was 99.9% LC-MS grade water (Thermo Fisher Scientific) with 0.1% formic acid and solvent “B” was 99.9% LC-MS grade acetonitrile (Thermo Fisher Scientific) with 0.1% formic acid. Peptides were loaded onto the trapping cartridge using a flow of 30 μL/min of solvent A for 3 min. Peptide elution was afterward performed with a constant flow of 0.3 μL/min using a total gradient time of 120 min. During the elution step, the percentage of solvent B was increased stepwise: 2% to 4% B in 4 min, from 4% to 8% in 2 min, 8% to 28% in 96 min, and from 28% to 40% in another 10 min. A column cleaning step using 80% B for 3 min was applied before the system was set again to its initial conditions (2% B) for re-equilibration for 10 minutes.

The peptides were introduced into the mass spectrometer Q Exactive Plus (Thermo Fisher Scientific) via a Pico-Tip Emitter 360 μm OD x 20 μm ID; 10 μm tip (New Objective). A spray voltage of 2.3 kV was applied and the mass spectrometer was operated in positive ion mode. The capillary temperature was 320°C. Full-scan MS spectra with a mass range of 375-1200 *m/z* were acquired in profile mode in the Orbitrap using a resolution of 70,000. The filling time was a maximum of 250 ms, and/or a maximum of 3e6 ions (automatic gain control, AGC) was collected. The instrument was cycling between MS and MS/MS acquisition in a data-dependent mode and consecutively fragmenting the Top 10 peaks of the MS scan. MS/MS spectra were acquired in profile mode in the Orbitrap with a resolution of 35,000, a maximum fill time of 120 ms and an AGC target of 2e5 ions. The quadrupole isolation window was set to 1.0 *m/z* and the first mass was fixed to 100 *m/z*. The normalized collision energy was 32 and the minimum AGC trigger was 2e2 ions (intensity threshold 1e3). Dynamic exclusion was applied and set to 30 s. The peptide match algorithm was set to ‘preferred’ and charge states ‘unassigned’, 1, 5 - 8 were excluded.

### Cells cryo plunge freezing and cryo-FIB milling

For vitrification of *E. coli* cells, 3.5 µl cell suspension were taken directly from the cell cultures and pipetted on Quantifoil R 1/1 grids glow discharged for 90 sec at 15 mA and 0.38 mbar using PELCO easiGlow. Grids were transferred to a Leica GP2 plunge freezer, excess liquid was blotted for 10 sec at 20 °C, grids were plunge frozen in liquid ethane and clipped into Autogrid sample holders. Autogrids were mounted into a FIB-shuttle and transferred using a cryo-transfer system to the cryo-stage of a dual-beam Aquilos FIB microscope (Thermo Scientific). Samples were coated with an organometallic platinum layer using a gas injection system for 10 sec and additionally sputter-coated with platinum at 1kV and 10 mA current for 10 sec. SEM imaging was performed at 2-10 kV and 13 pA. Milling was performed by a stepwise current decrease from 500 to 100 pA. Lamellae were polishing at 30-50 pA beam currents at shallow angles of 7-9°. Prior to grid unloading some lamellae were sputter-coated with platinum 1-2 sec at 1 kV and 10 mA. Grids were stored in liquid nitrogen upon data collection.

### Cryo-ET data collection

Cryo-ET data were collected on a Titan Krios G2 transmission electron microscope (Thermo Scientific) operated at 300 kV and equipped with a field-emission gun, a BioQuantum-K3 energy filter (Gatan). The K3 direct electron detector (Gatan) was operated in electron counting mode. Montages of individual lamellae were acquired at 6,500 X magnification with a 2.83 nm pixel size. Tilt series (TS) projections were acquired at a magnification of 53,000X, corresponding to a pixel size 1.697 Å at the specimen level over a tilt range of −63° to 45°, starting from an pre-tilt angle of 7-9 degrees to compensate for lamella pre-tilt relative to the grid, at 3° tilt increment, following a dose-symmetric scheme (Hagen et al., 2017), with tilt increments grouped by 2. The defocus range was set to −1.5 to −5 μm. total dose per tilt series to ∼ 120 e^-^/Å^2^, a 70 μm objective aperture was inserted and energy slit width was set to 20eV for TS collection. Tilt series images were acquired as 6K x 4K movies of 10 frames each, which were motion-corrected on-the-fly in SerialEM (Mastronarde, 2005). In total, 87 TS were acquired for *E. coli* K-12 LB, 82 TS for *E. coli* K-12 TET, 51 TS for *E. coli* ED1a LB, and 44 TS for *E. coli* ED1a TET samples. All of the cryo-ET data were collected for the timepoint of 30 min.

### Tilt series alignment and template matching

Motion-corrected tilt series were visually inspected to remove bad projections at high tilts. Then the TS were aligned through patch-tracking in IMOD (Kremer et al., 1996) and reconstructed using weighted back-projection with SIRT-like filtering of 10 iterations at a pixel size of 13.576 Å. These tomograms were used for visual inspection of tilt series alignment and tomogram thickness. Selected TS and their respective tomograms were reconstructed in Warp (Tegunov and Cramer, 2019) based on the IMOD-derived alignment files. Cross-correlation-based template matching in the Dynamo package (Castano-Diez et al., 2012) was used to localize ribosome particles within reconstructed tomograms. To prepare the *ab initio*-like reference, five tomograms generated in IMOD with SIRT-like filtering were matched using angular sampling of 30 degrees and 70S ribosome (EMDB-4050) as a template. The template was downsampled to 13.576 Å per voxel and filtered to ∼ 70 Å. The highest 1000 cross-correlation peaks were extracted as subtomograms at 6.788 Å pixel size. To avoid duplicates, after extraction of every particle, a 300 Å mask in diameter was applied to the cross-correlation volume. Subtomograms were aligned using NovaSTA package (Turoňová, Zenodo, 2022 https://github.com/turonova/novaSTA). The obtained structure was used as a reference for template matching of all the datasets in the Warp-generated tomograms. For consequent STA and classification, particles were extracted as described above. The particles list was converted to a Warp-compatible star-file using the *dynamo2m* toolbox (https://github.com/alisterburt/dynamo2m) (Burt et al., 2021).

### Subtomogram averaging and classification

The datasets corresponding to each of the four conditions, namely K-12/ED1a 30 min LB/TET were initially processed separately. The step-by-step processing scheme is described in **Figure S6**. Sub-tomograms were extracted with a pixel size of 6.788 Å and a box size of 64 pixels. The initial average was generated from the extracted subtomograms using the *relion_reconstruct* command and was used as a reference for the rough 3D classification in Relion 3.1 (Zivanov et al., 2018). The ribosomes were sorted from the membranes and junk particles using 3D classification with the following parameters: 7.5 degrees angular sampling, offset range 5 pixels, offset search step 1 pixel. To resolve the high-resolution subtomogram average of the *E. coli* 70S ribosome, the 70S particles from all datasets were merged and aligned in Relion and then refined using M (Tegunov et al., 2021) at a pixel size of 6.788 Å. The particles were extracted with 3.394 Å/px and a box size of 128 pixels, refined in Relion, and subjected to the focused 3D classification. The 75,670 well-aligned particles were re-extracted with 1.697 Å/px and a box size of 294 pixels and refined using 3 iterations in M to a final resolution of 5.3 Å.

To sample ribosomes states, 70S subtomograms were classified focusing on P-site tRNA and factor-binding regions without alignment. The resulting classes representing ribosomes at different states were refined in M. The following parameters were used for all 3D classification jobs: reference low-pass filter 50 Å, number of classes 3-6; T-parameter 10; 25-35 iterations; 350 Å mask diameter After every classification step, the selected classes were refined in Relion. The star files from Relion were converted to M-compatible format using the *relion_downgrade* command of the *dynamo2m* package.

### Code Availability

Script for measuring intermembrane distances is available at the repository https://github.com/martinschorb/membranedist.

## Acknowledgements

The authors would like to acknowledge Özkan Yildiz, Juan F. Castillo Hernandez, Thomas Hoffmann for support with scientific computing, Mark Linder and the Central Electron Microscopy facility of the Max Planck Institute of Biophysics for support with cryo-ET sample preparation and data acquisition. We thank Barbara Rathmann for assistance in MS data collection. The authors are greateful to Stefanie Böhm for critical reading and assesment of the manuscript. I.K. acknowledges Wim Hagen, and Felix Weiss for initial electron microscopy training; Matteo Allegretti, Maximilian Seidel, Patrick Hoffmann, and Andre Schwarz for discussions on the project. M.B. acknowledges funding by the Max Planck Society and EMBL.

## Author Contributions

I.K. conceived the project, designed experiments, performed experiments, analyzed data, and wrote the manuscript. N.R. developed the TPP analysis pipeline, analyzed data, and wrote the manuscript. C.G. and A.T. conceived the project, advised and provided protocols for microbiology experiments, edited the manuscript. I.L.I performed polysome analysis and assisted computational data analysis under supervision of F.J.T., B.T. and C.E.Z. supported cryo-ET data processing. S.W assisted cryo-ET data acquisition. J.D.L. supervised mass spectromerty experiments, M.B. conceived the project, supervised the project, analyzed data, acquired funding, and wrote the manuscript.

## Accession Numbers

Cryo-ET density maps generated in this study have been deposited in the EM Data Bank with the following accession codes: EMD-AAAAA (in situ *E. coli* 70S ribosome), EMD-BBBBB (in situ 70S ribosome of *E. coli* K-12 untreated cells), EMD-CCCCC (in situ 70S ribosome of *E. coli* K-12 cells treated with tetracycline), EMD-DDDDD (In situ 70S ribosome of *E. coli* ED1a untreated cells), EMD-EEEEE (in situ 70S ribosome of *E. coli* ED1a cells treated with tetracycline), EMD-FFFFF (*E. coli* K-12 70S ribosome bound to mRNA A-tRNA, P-tRNA, E-tRNA), EMD-GGGGG (*E. coli* ED1a 70S ribosome bound to mRNA A-tRNA, P-tRNA, E-tRNA). All proteomics data associated with this manuscript have been uploaded to the PRIDE online repository.

---

**Figure S1.**
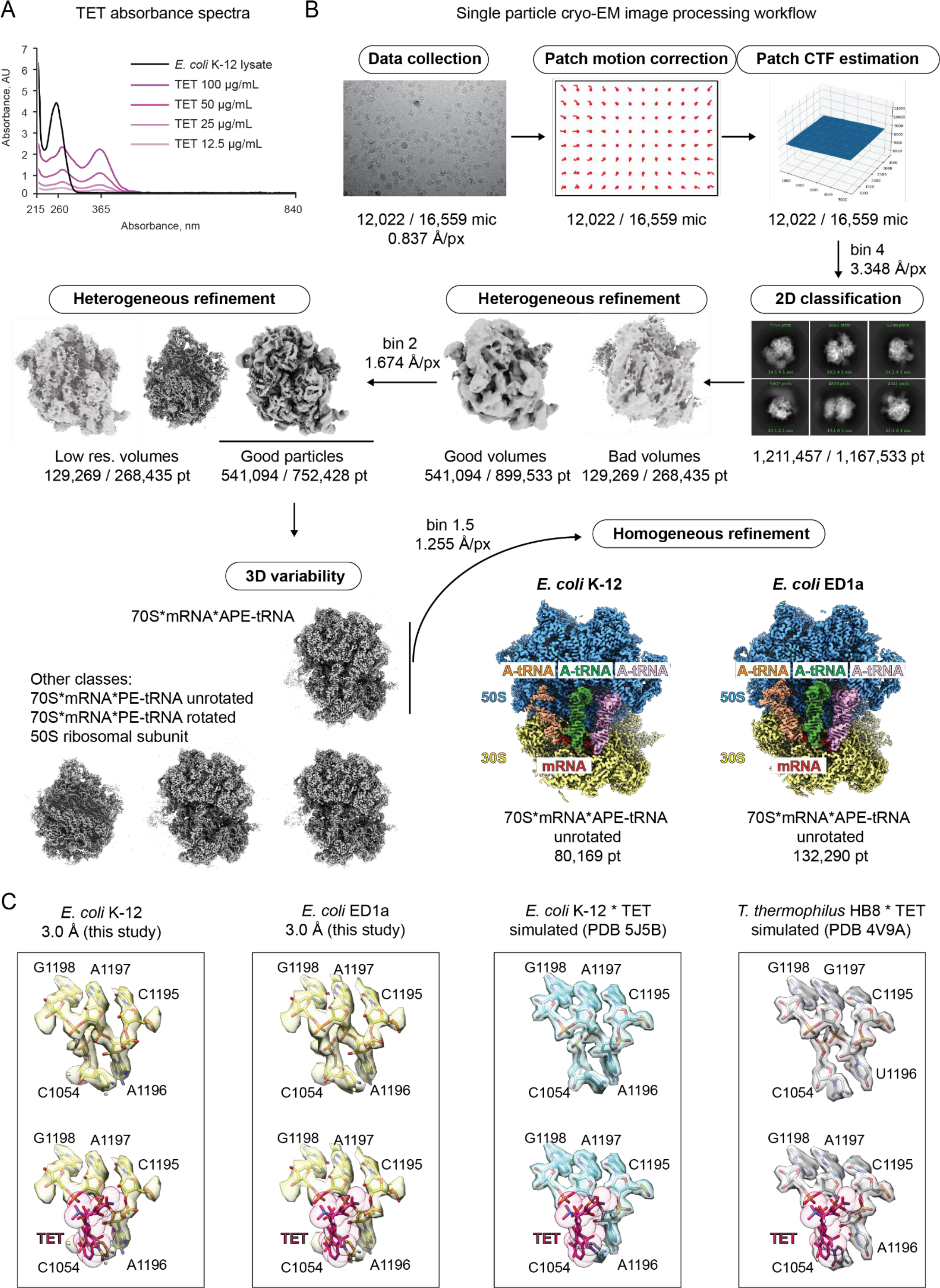
Structural analysis of the ribosomes reveals identical structure of TET-binding pocket in bacteria. (A) The absorbance spectrum of the cell lysate (black) and TET is titrated in phosphate buffer saline to concentrations ranging from 12.5 to 100 μg/ml (shades of purple). TET has an absorption peak at 365 nm, which is not present in the untreated cell lysate. (B) Classification strategy used to obtain 70S structures using single particle cryo-EM. For simplicity, the intermediate reconstructions (gray) are shown only for *E. coli* K-12 dataset. The numbers denoted below the images correspond to the micrographs (mic) and particles (pt) analyzed at each step for the K-12 strain and ED1a, respectively. The final reconstructions are shown for both strains. (C) Comparison of the TET binding sites of the *E. coli* K-12 and *E. coli* ED1a 70S·mRNA·APE-tRNAs structures from the current study (yellow) with published crystal structures of the 70S·TET complex from *E. coli* K-12 (cyan, (Cocozaki et al., 2016)) and *T. thermophilus* (gray, (Jenner et al., 2013). The TET binding site of crystal structures was simulated at 3.0 Å around the selected nucleotides using the *molmap* function in ChimeraX. The upper image in each panel shows the pocket without TET, the lower image includes antibiotic (magenta).

**Figure S2.**
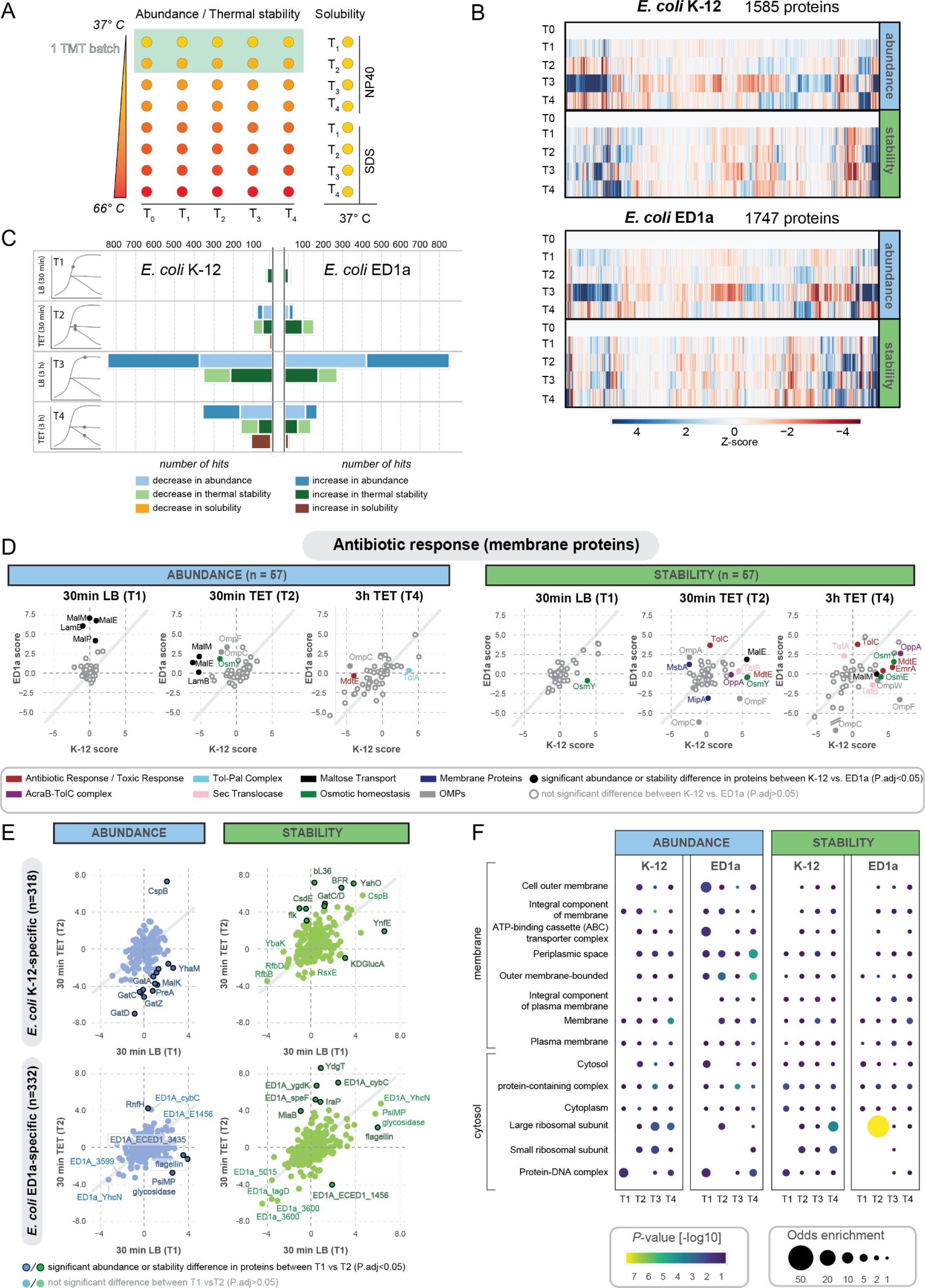
2D-TPP identifies changes in abundance and thermal stability for proteins involved in antibiotic response and several uncharacterized proteins. (A) Schematic setup of 2D-TPP analysis applied in this study. For abundance and thermal stability measurements (left), protein fractions were used upon heating from 37 °C to 66 °C. For solubility measurements, the experiments were done in the presence of NP40 or SDS, respectively, at a constant temperature of 37 °C. Each sample was prepared in four biological replicates. (B) Heatmap representation of abundance (upper block), and thermal stability (lower block) changes of proteins identified in the TPP screen. Coloring corresponds to the calculated z-score ranging from −4 (red) to +4 (blue). The timepoint T3 (3 h LB) shows drastic proteome rearrangements, likely due to a shift to stationary phase of both strains. (C) Barplot illustrating the number of significantly affected proteins in abundance (blue shades), thermal stability (green shades), and solubility (orange/brown shades) for *E. coli* K-12 (left) and *E. coli* ED1a (right) for all analyzed timepoints. Timepoints T1 and T3 represent untreated cells grown for 30 min and 3 h, respectively. Timepoints T2 and T4 represent TET treated cells grown for 30 min and 3 h, respectively. The scores are calculated relative to the control condition T0 (untreated cells, 0 min LB). (D) Scatter plot of abundance (left) and stability values (right) for annotated membrane proteins involved in antibiotic response according to gene onthology mapping for T1 (30 min LB), T2 (30 min TET) and T4 (3 h TET), respectively, comparing K-12 scores (*x*-axis) against ED1a scores (*y*-axis). Each dot represents a protein; if it is highlighted in color, it signifies that there is a significant difference between K-12 and ED1a scores (*P_adj_*<0.05). The color scheme is explained in the legend below. (E) Scatter plot of abundance (left) and stability values (right) for protein hits detected only in K-12 (upper panels) or ED1a (lower panels) samples, comparing T1 (30 min LB) scores (*x*-axis) against T2 (30 min TET) scores (*y*-axis). Each dot represents a protein; highlighted black contours indicate a significant difference between T1 and T2 scores (*P_adj_*<0.05). The color scheme is explained in the legend below the figure panel. (F) Gene ontology (GO) enrichment plot for cellular compartments, with compartments ordered according to their physical location on the outer membrane – cytosol axis for all analyzed timepoints. The enrichment for each GO-term in the significant protein set (global & local FDR < 0.05) in each respective condition (*x*-axis) is calculated using the Fisher Exact Test, relative to the insignificant portion of proteins in each condition. Bubbles are displayed for GO-terms that are enriched in at least one condition (*P*<0.01) and have at least 10 protein components significantly affected in the dataset,. The bubble size indicates the odds enrichment, whereas the bubble color reflects the *P*-value (-log10 scale).

**Figure S3.**
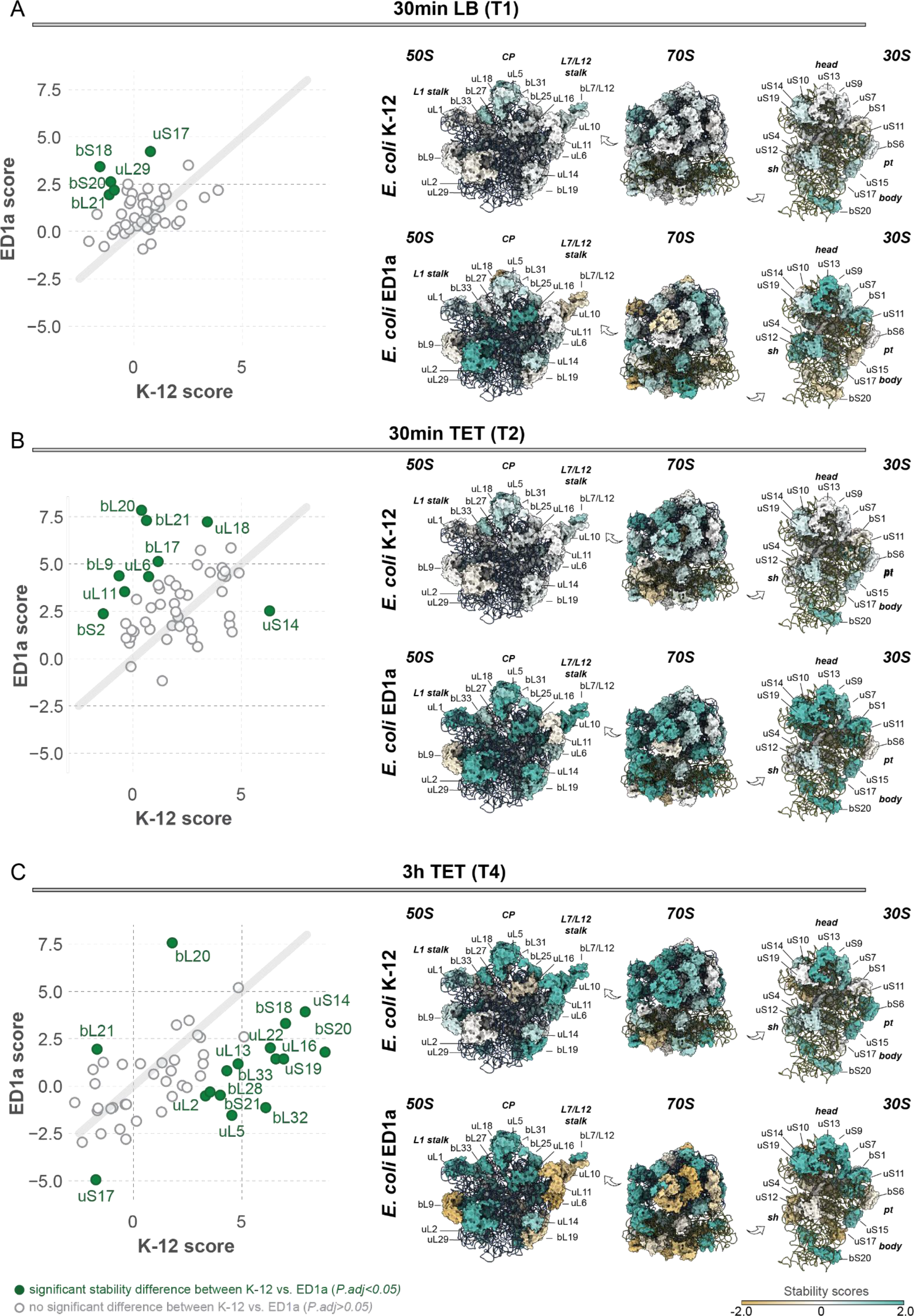
TPP values mapped on ribosome structure show thermal stability increase for interface r-proteins and decrease for peripheral r-proteins in *E. coli* ED1a upon TET treatment. (A-C) Left-hand side: scatter plot of stability values for r-proteins for (A) T1 (30 min LB), (B) T2 (30 min TET), (C) for T4 (3 h TET) comparing K-12 *Z*-scores (*x*-axis) against ED1a scores (*y*-axis). Each dot represents a protein; significant differences between K-12 and ED1a scores are coloredgreen (*P_adj_*<0.05). Right-hand side: structure of the 50S, 70S, and 30S ribosome structures in K-12 (upper) and ED1a (lower), shaded according to the respective thermal stability score values in *E. coli* K-12 and ED1a, respectively. The PDB model was combined from coordinates fetched from PDB 3J7Z, PDB 7K00, and PDB 6H4N. Proteins with increased stability are colored turquoise, proteins with decreased stability are colored tan, and proteins with unchanged stability are colored white.

**Figure S4.**
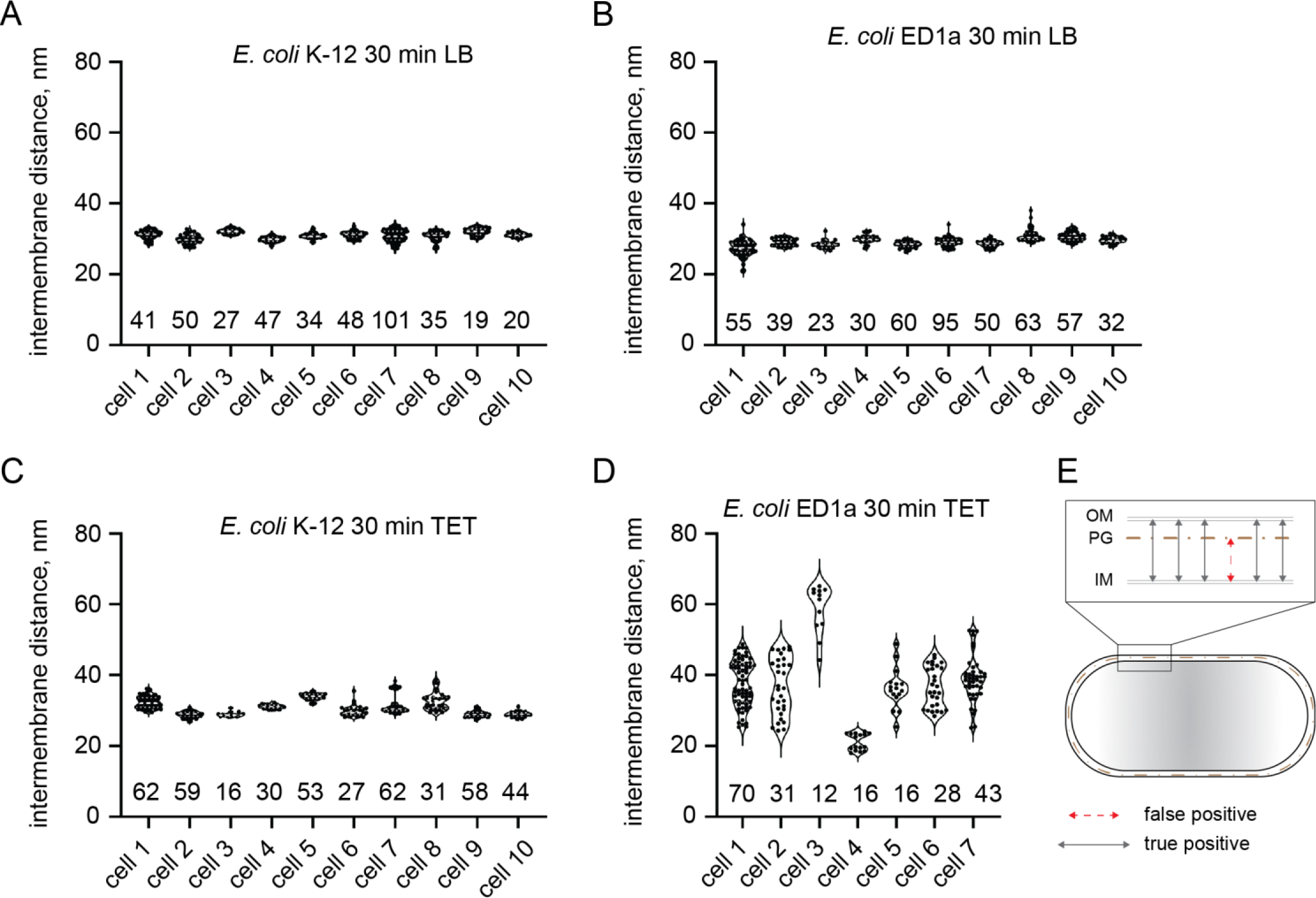
Intermembrane distance measurements of cells used for subtomogram averaging indicate membrane wobbling in each individual cell of the ED1a-TET sample. (A) – (D) Measurements of intermembrane distances in individual cells from Figure 4B. For calculation details, refer to Figure 4B in the main text. Overall, the thickness of periplasmic space for samples K-12 LB (A), ED1a LB (B), and K-12 TET (C) was constant within each cell and varied slightly between the cells. For *E. coli* ED1 TET sample (D), the periplasmic space varied both within single cells and among cells. (E) Schematic representation of intermembrane distance measurement. OM – outer membrane, IM – inner membrane, PG – peptydoglycal layer.

**Figure S5.**
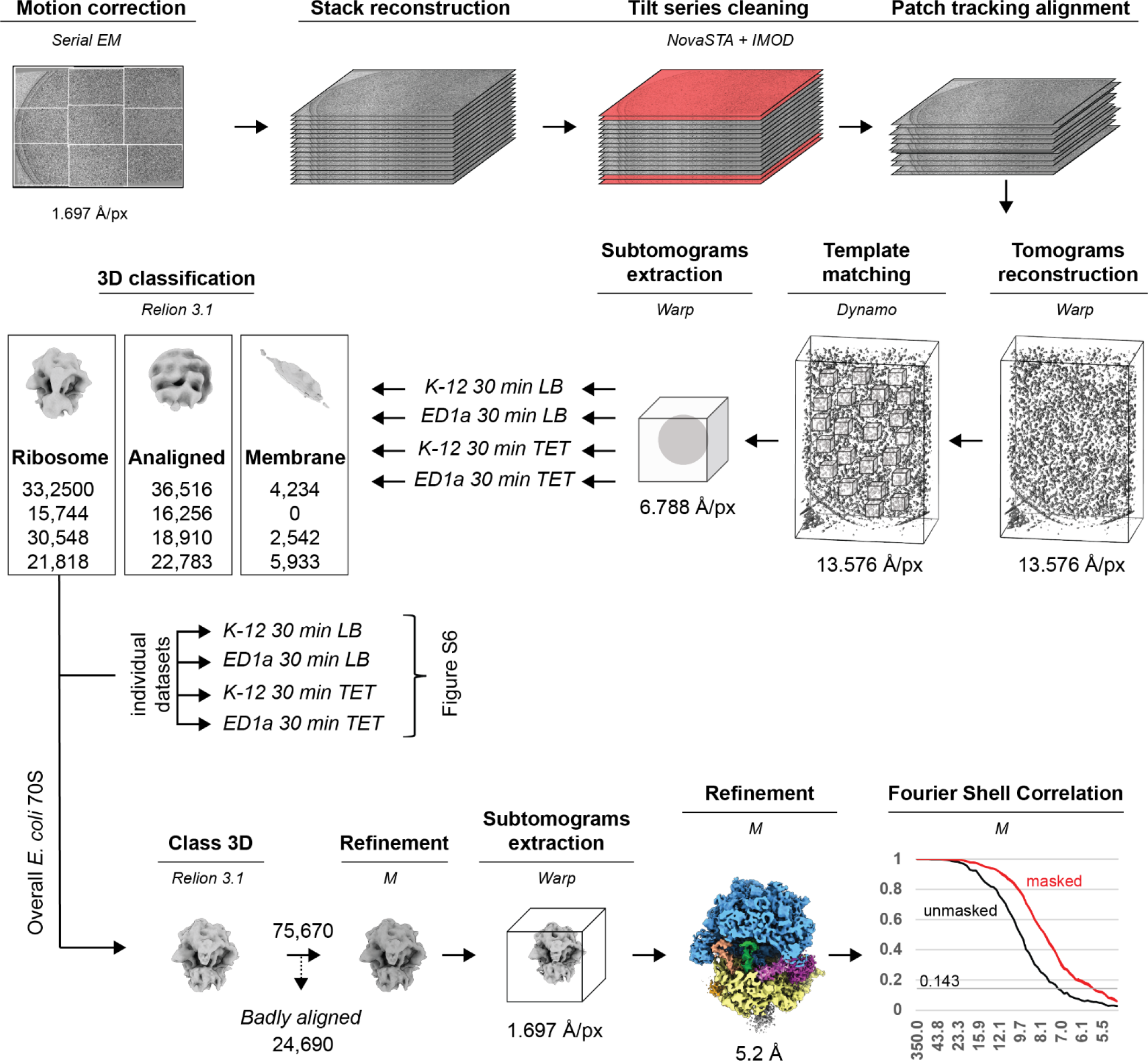
Applied cryo-ET image processing pipeline achieves high-resolution 70S ribosome structure. Image processing of cryo-ET data. Each step is highlighted in bold letters, and software used is denoted in italics. The number of particles is indicated below each density snapshot. The pixel size values reflect data binning and un-binning. All ribosomal subtomograms were merged and aligned in Relion 3.1 and refined in M to an overall structure of 5.2 Å resolution. The detailed scheme of 3D classification used to sort out the structures of 70S with different or ligand occupancies in individual datasets is shown in Figure S2.

**Figure S6.**
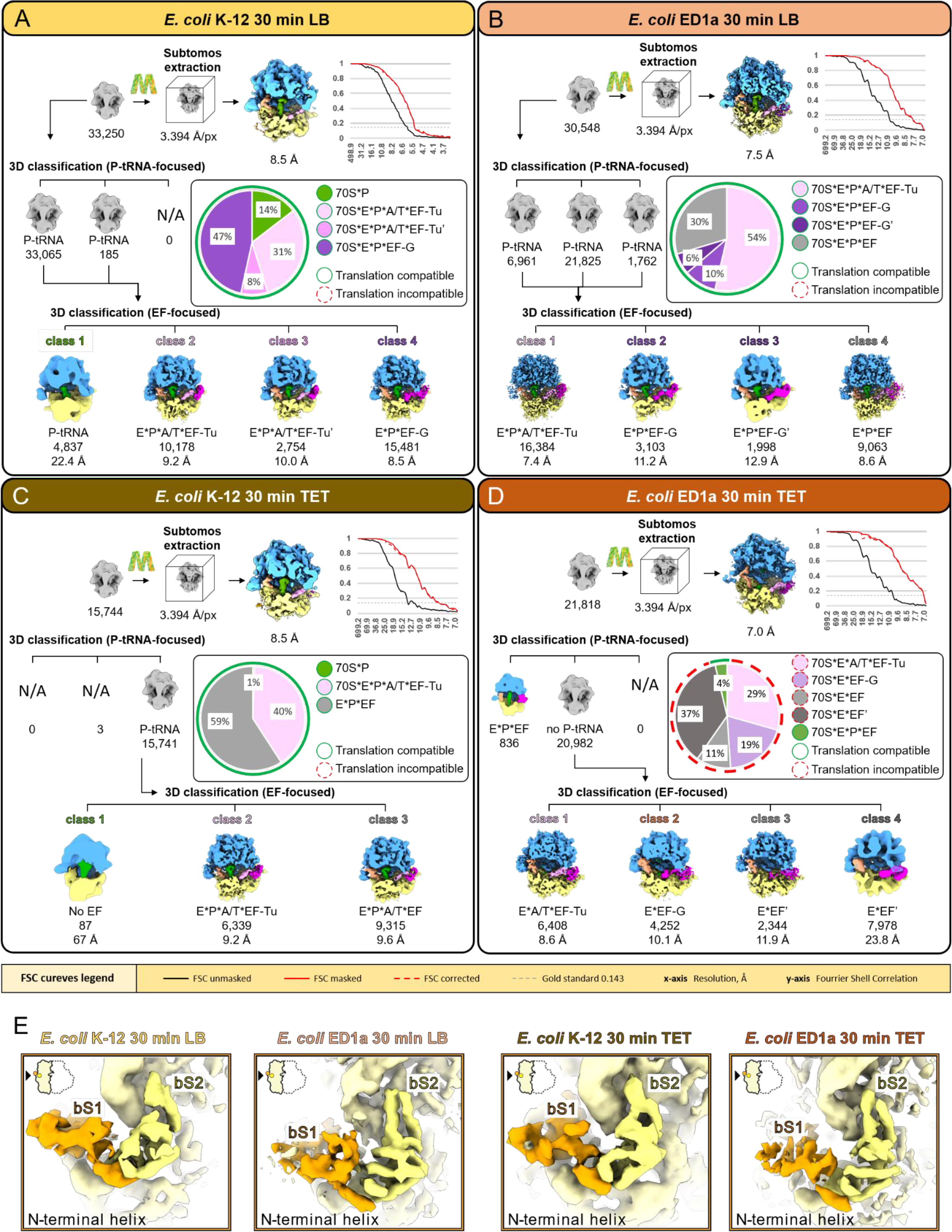
Subtomogram averaging and 3D classification reveals that the majority of ribosomes are present in a translation-incompetent state in *E. coli* ED1a cells after 30 min of TET treatment. (A) – (D) Multiple rounds of 3D classification allow identification of 70S structures with different ligand occupancies. The distribution of classes in every sample is shown as a pie chart. The classes containing or lacking tRNA in the P-site are highlighted around the pie charts as green and red circles, respectively. The classes with ambiguous density at the elongation factor binding site are captioned as EF. (E) Segmented density of the r-protein bS1 (orange) anchored with its N-terminal helix to the r-protein bS2 (yellow) on the small ribosomal subunits of 70S averages from *E. coli* K-12 30 min LB, *E. coli* K-12 30 min TET, *E. coli* ED1a 30 min LB, *E. coli* ED1a 30 min TET.

**Table S1.** Thermal proteome profiling hits (External file Excel sheet). Related to STAR Methods.

**Table S2.** Single particle cryo-EM data collection and data processing statistics. Related to STAR Methods.

|  | *E. coli* K-12 30 min LB<br>(EMDB-XXXX) | *E. coli* K-12 30 min TET<br>(EMDB-YYYY) |
| --- | --- | --- |
| Magnification | 105,000 X | 105,000 X |
| Voltage (kV) | 300 | 300 |
| Exposure dose (e-/Å<sup>2</sup>) | 40 | 40 |
| Number of frames | 40 | 40 |
| Defocus range (μm) | -0.25 to -2.5 | -0.25 to -2.5 |
| Pixel size (Å/px) | 0.837 | 0.837 |
| No. of micrographs | 12,022 | 16,559 |
| No. of extracted particles | 1,740,048 | 2,302,674 |
| Final no. of particles | 80,169 | 132,290 |
| FSC threshold | 0.143 | 0.143 |
| Map resolution (Å) | 3.0 | 3.0 |

**Table S3.** Cryo-ET data collection and sub-tomogram averaging statistics. Related to STAR Methods.

| Parameter | *E. coli*<br>combined<br>(EMDB-AAAAA) | *E. coli* K-12<br>30 min LB<br>(EMDB-BBBBB) | *E. coli* K-12<br>30 min TET<br>(EMDB-BBBBB) | *E. coli* ED1a<br>30 min LB<br>(EMDB-CCCCC) | *E. coli* ED1a<br>30 min TET<br>(EMDB-DDDDD) |
| --- | --- | --- | --- | --- | --- |
| **Data collection and processing** |  |  |  |  |  |
| Magnification | 53,000 X | 53,000 X | 53,000 X | 53,000 X | 53,000 X |
| Voltage (kV) | 300 | 300 | 300 | 300 | 300 |
| Exposure dose (e-/Å<sup>2</sup>) | 120-130 | 120-130 | 120-130 | 120-130 | 120-130 |
| Defocus range (μm) | -1.5 to -4.5 | -1.5 to -4.5 | -1.5 to -3.5 | -1.5- to 4.0 | -1.5 to -3.5 |
| Pixel size (Å/px) | 1.697 | 1.697 | 1.697 | 1.697 | 1.697 |
| No. of collected tilt series | 264 | 87 | 82 | 51 | 44 |
| No. of used tomograms | 104 | 34 | 17 | 26 | 27 |
| No. of extracted particles | 208,000 | 68,000 | 34,000 | 52,000 | 54,000 |
| Final no. of particles | 75,670 | 33,250 | 15,744 | 30,548 | 21,818 |
| FSC threshold | 0.143 | 0.143 | 0.143 | 0.143 | 0.143 |
| Map resolution (Å) | 5.3 | 8.5 | 8.5 | 7.5 | 7.0 |

## References

1. Baba, T., Ara, T., Hasegawa, M., Takai, Y., Okumura, Y., Baba, M., Datsenko, K.A., Tomita, M., Wanner, B.L., and Mori, H. (2006). Construction of Escherichia coli K-12 in-frame, single-gene knockout mutants: the Keio collection. Mol Syst Biol 2, 2006 0008.

2. Becher, I., Andres-Pons, A., Romanov, N., Stein, F., Schramm, M., Baudin, F., Helm, D., Kurzawa, N., Mateus, A., Mackmull, M.T., et al. (2018). Pervasive Protein Thermal Stability Variation during the Cell Cycle. Cell 173, 1495–1507 e1418.

3. Beckert, B., Turk, M., Czech, A., Berninghausen, O., Beckmann, R., Ignatova, Z., Plitzko, J.M., and Wilson, D.N. (2018). Structure of a hibernating 100S ribosome reveals an inactive conformation of the ribosomal protein S1. Nat Microbiol 3, 1115–1121.

4. Brodersen, D.E., Clemons, W.M., Jr., Carter, A.P., Morgan-Warren, R.J., Wimberly, B.T., and Ramakrishnan, V. (2000). The structural basis for the action of the antibiotics tetracycline, pactamycin, and hygromycin B on the 30S ribosomal subunit. Cell 103, 1143–1154.

5. Burt, A., Gaifas, L., Dendooven, T., and Gutsche, I. (2021). A flexible framework for multi-particle refinement in cryo-electron tomography. Plos Biol 19, e3001319.

6. Castano-Diez, D., Kudryashev, M., Arheit, M., and Stahlberg, H. (2012). Dynamo: a flexible, user-friendly development tool for subtomogram averaging of cryo-EM data in high-performance computing environments. J Struct Biol 178, 139–151.

7. Cheng-Guang, H., and Gualerzi, C.O. (2021). The Ribosome as a Switchboard for Bacterial Stress Response. Frontiers in Microbiology 11.

8. Clermont, O., Lescat, M., O’Brien, C.L., Gordon, D.M., Tenaillon, O., and Denamur, E. (2008). Evidence for a human-specific Escherichia coli clone. Environ Microbiol 10, 1000–1006.

9. Cocozaki, A.I., Altman, R.B., Huang, J., Buurman, E.T., Kazmirski, S.L., Doig, P., Prince, D.B., Blanchard, S.C., Cate, J.H., and Ferguson, A.D. (2016). Resistance mutations generate divergent antibiotic susceptibility profiles against translation inhibitors. Proc Natl Acad Sci U S A 113, 8188–8193.

10. Demo, G., Rasouly, A., Vasilyev, N., Svetlov, V., Loveland, A.B., Diaz-Avalos, R., Grigorieff, N., Nudler, E., and Korostelev, A.A. (2017). Structure of RNA polymerase bound to ribosomal 30S subunit. Elife 6.

11. Duval, M., Korepanov, A., Fuchsbauer, O., Fechter, P., Haller, A., Fabbretti, A., Choulier, L., Micura, R., Klaholz, B.P., Romby, P., et al. (2013). Escherichia coli ribosomal protein S1 unfolds structured mRNAs onto the ribosome for active translation initiation. Plos Biol 11, e1001731.

12. Emsley, P., and Cowtan, K. (2004). Coot: model-building tools for molecular graphics. Acta Crystallogr D Biol Crystallogr 60, 2126–2132.

13. Fedry, J., Silva, J., Vanevic, M., Fronik, S., Mechulam, Y., Schmitt, E., Georges, A.d., Faller, W., and Förster, F. (2023). Visualization of translation reorganization upon persistent collision stress in mammalian cells. 2023.2003.2023.533914.

14. Gemmer, M., Chaillet, M.L., van Loenhout, J., Cuevas Arenas, R., Vismpas, D., Grollers-Mulderij, M., Koh, F.A., Albanese, P., Scheltema, R.A., Howes, S.C., et al. (2023). Visualization of translation and protein biogenesis at the ER membrane. Nature 614, 160–167.

15. Gilbert, J.A., Blaser, M.J., Caporaso, J.G., Jansson, J.K., Lynch, S.V., and Knight, R. (2018). Current understanding of the human microbiome. Nat Med 24, 392–400.

16. Gromadski, K.B., and Rodnina, M.V. (2004). Kinetic determinants of high-fidelity tRNA discrimination on the ribosome. Mol Cell 13, 191–200.

17. Hagen, W.J.H., Wan, W., and Briggs, J.A.G. (2017). Implementation of a cryo-electron tomography tilt-scheme optimized for high resolution subtomogram averaging. J Struct Biol 197, 191–198.

18. Hoffmann, P.C., Kreysing, J.P., Khusainov, I., Tuijtel, M.W., Welsch, S., and Beck, M. (2022). Structures of the eukaryotic ribosome and its translational states in situ. Nat Commun 13, 7435.

19. Jenner, L., Starosta, A.L., Terry, D.S., Mikolajka, A., Filonava, L., Yusupov, M., Blanchard, S.C., Wilson, D.N., and Yusupova, G. (2013). Structural basis for potent inhibitory activity of the antibiotic tigecycline during protein synthesis. Proc Natl Acad Sci U S A 110, 3812–3816.

20. Jones-Dias, D., Carvalho, A.S., Moura, I.B., Manageiro, V., Igrejas, G., Canica, M., and Matthiesen, R. (2017). Quantitative proteome analysis of an antibiotic resistant Escherichia coli exposed to tetracycline reveals multiple affected metabolic and peptidoglycan processes. J Proteomics 156, 20–28.

21. Khanna, K., and Villa, E. (2022). Revealing bacterial cell biology using cryo-electron tomography. Curr Opin Struct Biol 75, 102419.

22. Khusainov, I., Fatkhullin, B., Pellegrino, S., Bikmullin, A., Liu, W.T., Gabdulkhakov, A., Shebel, A.A., Golubev, A., Zeyer, D., Trachtmann, N., et al. (2020). Mechanism of ribosome shutdown by RsfS in Staphylococcus aureus revealed by integrative structural biology approach. Nat Commun 11, 1656.

23. Khusainov, I., Vicens, Q., Ayupov, R., Usachev, K., Myasnikov, A., Simonetti, A., Validov, S., Kieffer, B., Yusupova, G., Yusupov, M., et al. (2017). Structures and dynamics of hibernating ribosomes from Staphylococcus aureus mediated by intermolecular interactions of HPF. Embo J 36, 2073–2087.

24. Khusainov, I., Vicens, Q., Bochler, A., Grosse, F., Myasnikov, A., Menetret, J.F., Chicher, J., Marzi, S., Romby, P., Yusupova, G., et al. (2016). Structure of the 70S ribosome from human pathogen Staphylococcus aureus. Nucleic Acids Res 44, 10491–10504.

25. Kremer, J.R., Mastronarde, D.N., and McIntosh, J.R. (1996). Computer visualization of three-dimensional image data using IMOD. J Struct Biol 116, 71–76.

26. Levin, B.R., McCall, I.C., Perrot, V., Weiss, H., Ovesepian, A., and Baquero, F. (2017). A Numbers Game: Ribosome Densities, Bacterial Growth, and Antibiotic-Mediated Stasis and Death. Mbio 8.

27. Maier, L., Goemans, C.V., Wirbel, J., Kuhn, M., Eberl, C., Pruteanu, M., Muller, P., Garcia-Santamarina, S., Cacace, E., Zhang, B., et al. (2021). Unravelling the collateral damage of antibiotics on gut bacteria. Nature 599, 120–124.

28. Mastronarde, D.N. (2005). Automated electron microscope tomography using robust prediction of specimen movements. J Struct Biol 152, 36–51.

29. O’Reilly, F.J., Xue, L., Graziadei, A., Sinn, L., Lenz, S., Tegunov, D., Blotz, C., Singh, N., Hagen, W.J.H., Cramer, P., et al. (2020). In-cell architecture of an actively transcribing-translating expressome. Science 369, 554–557.

30. Oliva, B., Gordon, G., McNicholas, P., Ellestad, G., and Chopra, I. (1992). Evidence that tetracycline analogs whose primary target is not the bacterial ribosome cause lysis of Escherichia coli. Antimicrob Agents Chemother 36, 913–919.

31. Ortiz, J.O., Etchells, S., Leis, A., Hartl, F.U., and Baumeister, W. (2007). Ribosomes under stress: Structural variability of the 100S particles studied by cryoelectron tomography. J Biomol Struct Dyn 24, 633–633.

32. Pettersen, E.F., Goddard, T.D., Huang, C.C., Meng, E.C., Couch, G.S., Croll, T.I., Morris, J.H., and Ferrin, T.E. (2021). UCSF ChimeraX: Structure visualization for researchers, educators, and developers. Protein Sci 30, 70–82.

33. Polikanov, Y.S., Blaha, G.M., and Steitz, T.A. (2012). How hibernation factors RMF, HPF, and YfiA turn off protein synthesis. Science 336, 915–918.

34. Prossliner, T., Skovbo Winther, K., Sorensen, M.A., and Gerdes, K. (2018). Ribosome Hibernation. Annu Rev Genet 52, 321–348.

35. Punjani, A., Rubinstein, J.L., Fleet, D.J., and Brubaker, M.A. (2017). cryoSPARC: algorithms for rapid unsupervised cryo-EM structure determination. Nat Methods 14, 290–296.

36. Qu, X., Lancaster, L., Noller, H.F., Bustamante, C., and Tinoco, I., Jr. (2012). Ribosomal protein S1 unwinds double-stranded RNA in multiple steps. Proc Natl Acad Sci U S A 109, 14458–14463.

37. Rodnina, M.V. (2018). Translation in Prokaryotes. Cold Spring Harb Perspect Biol 10.

38. Savelsbergh, A., Katunin, V.I., Mohr, D., Peske, F., Rodnina, M.V., and Wintermeyer, W. (2003). An elongation factor G-induced ribosome rearrangement precedes tRNA-mRNA translocation. Mol Cell 11, 1517–1523.

39. Savitski, M.M., Reinhard, F.B., Franken, H., Werner, T., Savitski, M.F., Eberhard, D., Martinez Molina, D., Jafari, R., Dovega, R.B., Klaeger, S., et al. (2014). Tracking cancer drugs in living cells by thermal profiling of the proteome. Science 346, 1255784.

40. Sorensen, M.A., Fricke, J., and Pedersen, S. (1998). Ribosomal protein S1 is required for translation of most, if not all, natural mRNAs in Escherichia coli in vivo. J Mol Biol 280, 561–569.

41. Tegunov, D., and Cramer, P. (2019). Real-time cryo-electron microscopy data preprocessing with Warp. Nat Methods 16, 1146–1152.

42. Tegunov, D., Xue, L., Dienemann, C., Cramer, P., and Mahamid, J. (2021). Multi-particle cryo-EM refinement with M visualizes ribosome-antibiotic complex at 3.5 A in cells. Nat Methods 18, 186–193.

43. B. Turoňová, turonova/novaSTA: Advanced particle analysis, version 1.1, Zenodo (2022); https://doi.org/10.5281/zenodo.5921012.

44. VanBogelen, R.A., and Neidhardt, F.C. (1990). Ribosomes as sensors of heat and cold shock in Escherichia coli. Proc Natl Acad Sci U S A 87, 5589–5593.

45. Voorhees, R.M., and Ramakrishnan, V. (2013). Structural basis of the translational elongation cycle. Annu Rev Biochem 82, 203–236.

46. Wenzel, M., Dekker, M.P., Wang, B., Burggraaf, M.J., Bitter, W., van Weering, J.R.T., and Hamoen, L.W. (2021). A flat embedding method for transmission electron microscopy reveals an unknown mechanism of tetracycline. Commun Biol 4, 306.

47. Xing, H., Taniguchi, R., Khusainov, I., Kreysing, J.P., Welsch, S., Turoňová, B., and Beck, M. (2023). Translation dynamics in human cells visualized at high-resolution reveal cancer drug action. 2023.2003.2002.529652.

48. Xu, C., Lin, X., Ren, H., Zhang, Y., Wang, S., and Peng, X. (2006). Analysis of outer membrane proteome of Escherichia coli related to resistance to ampicillin and tetracycline. Proteomics 6, 462–473.

49. Xue, L., Lenz, S., Zimmermann-Kogadeeva, M., Tegunov, D., Cramer, P., Bork, P., Rappsilber, J., and Mahamid, J. (2022). Visualizing translation dynamics at atomic detail inside a bacterial cell. Nature 610, 205–211.

50. Yoshida, H., and Wada, A. (2014). The 100S ribosome: ribosomal hibernation induced by stress. Wiley Interdiscip Rev RNA 5, 723–732.

51. Yun, S.H., Choi, C.W., Park, S.H., Lee, J.C., Leem, S.H., Choi, J.S., Kim, S., and Kim, S.I. (2008). Proteomic analysis of outer membrane proteins from Acinetobacter baumannii DU202 in tetracycline stress condition. J Microbiol 46, 720–727.

52. Zhang, D.F., Jiang, B., Xiang, Z.M., and Wang, S.Y. (2008). Functional characterisation of altered outer membrane proteins for tetracycline resistance in Escherichia coli. Int J Antimicrob Agents 32, 315–319.

53. Zietek, M., Miguel, A., Khusainov, I., Shi, H., Asmar, A.T., Ram, S., Wartel, M., Sueki, A., Schorb, M., Goulian, M., et al. (2022). Bacterial cell widening alters periplasmic size and activates envelope stress responses. BioRxiv.

54. Zivanov, J., Nakane, T., Forsberg, B.O., Kimanius, D., Hagen, W.J., Lindahl, E., and Scheres, S.H. (2018). New tools for automated high-resolution cryo-EM structure determination in RELION-3. Elife 7.

## SUPPLEMENTAL REFERENCES

1. Brodersen, D.E., Clemons, W.M., Jr., Carter, A.P., Morgan-Warren, R.J., Wimberly, B.T., and Ramakrishnan, V. (2000). The structural basis for the action of the antibiotics tetracycline, pactamycin, and hygromycin B on the 30S ribosomal subunit. Cell 103, 1143–1154.

2. Cocozaki, A.I., Altman, R.B., Huang, J., Buurman, E.T., Kazmirski, S.L., Doig, P., Prince, D.B., Blanchard, S.C., Cate, J.H., and Ferguson, A.D. (2016). Resistance mutations generate divergent antibiotic susceptibility profiles against translation inhibitors. Proc Natl Acad Sci U S A 113, 8188–8193.

3. Jenner, L., Starosta, A.L., Terry, D.S., Mikolajka, A., Filonava, L., Yusupov, M., Blanchard, S.C., Wilson, D.N., and Yusupova, G. (2013). Structural basis for potent inhibitory activity of the antibiotic tigecycline during protein synthesis. Proc Natl Acad Sci U S A 110, 3812–3816.

4. Liu, Y.T., Zhang, H., Wang, H., Tao, C.L., Bi, G.Q., and Zhou, Z.H. (2022). Isotropic reconstruction for electron tomography with deep learning. Nat Commun 13, 6482.

